# In vivo reduction of age-dependent neuromelanin accumulation mitigates features of Parkinson’s disease

**DOI:** 10.1101/2022.08.08.503142

**Authors:** Marta Gonzalez-Sepulveda, Joan Compte, Thais Cuadros, Alba Nicolau, Camille Guillard-Sirieix, Núria Peñuelas, Marina Lorente-Picon, Annabelle Parent, Jordi Romero-Giménez, Joana M. Cladera-Sastre, Ariadna Laguna, Miquel Vila

## Abstract

Humans accumulate with age the dark-brown pigment neuromelanin inside specific neuronal groups. Neurons with the highest neuromelanin levels are particularly susceptible to degeneration in Parkinson’s disease, especially dopaminergic neurons of the *substantia nigra* (SN), the loss of which leads to characteristic motor Parkinson’s disease symptoms. In contrast to humans, neuromelanin does not appear spontaneously in most animals, including rodents, and Parkinson’s disease is an exclusively human condition. Using humanized neuromelanin-producing rodents, we recently found that neuromelanin can trigger Parkinson’s disease pathology when accumulated above a specific pathogenic threshold.

Here, by taking advantage of this newly developed animal model, we assessed whether the intracellular buildup of neuromelanin that occurs with age can be slowed down *in vivo* to prevent or attenuate Parkinson’s disease. Because neuromelanin derives from the oxidation of free cytosolic dopamine, we enhanced dopamine vesicular encapsulation in the SN of neuromelanin-producing rats by viral vector-mediated overexpression of vesicular monoamine transporter 2 (VMAT2). This strategy reduced the formation of potentially toxic oxidized dopamine species that can convert into neuromelanin and maintained intracellular neuromelanin levels below their pathogenic threshold. Decreased neuromelanin production was associated with an attenuation of Lewy body-like inclusion formation and a long-term preservation of dopamine homeostasis, nigrostriatal neuronal integrity and motor function in these animals.

Our results demonstrate the feasibility and therapeutic potential of modulating age-dependent intracellular neuromelanin production *in vivo*, thereby opening an unexplored path for the treatment of Parkinson’s disease and, in a broader sense, brain aging.

## Introduction

In Parkinson’s disease there is a preferential degeneration of neurons that contain the pigment neuromelanin (NM), especially dopamine (DA)-producing neurons of the *substantia nigra* pars compacta (SNpc), the loss of which leads to a depletion of nigral and striatal DA that underlies classical Parkinson’s disease motor symptoms. NM is a synthetic byproduct of DA metabolism that progressively accumulates with age, with neurons reaching the highest levels of NM being the most susceptible to Parkinson’s disease degeneration^1–3^. In contrast to humans, NM does not appear spontaneously in most animal species, including rodents^4^. However, we recently developed the first experimental rodent model exhibiting age-dependent production and accumulation of NM within Parkinson’s disease-vulnerable DA nigral neurons, at levels up to those reached in elderly humans. This model is based on the viral vector-mediated expression of melanin-producing enzyme tyrosinase (TYR) in rat SNpc^5^. Using this humanized NM-producing animal model, we revealed that age-dependent progressive intracellular build-up of NM ultimately compromised neuronal function and viability when allowed to accumulate above a specific threshold. As a consequence, these animals eventually developed major Parkinson’s disease -like features, including motor deficits, Lewy body (LB)- like pathology, nigrostriatal neurodegeneration and neuroinflammatory changes^5^. Relevant to humans, intracellular NM levels reach this pathogenic threshold in Parkinson’s disease patients and pre-Parkinson’s disease subjects^5^. These results indicate that an excessive production of NM within neurons can compromise neuronal function and trigger Parkinson’s disease pathology. Accordingly, strategies to slow down age-dependent NM production/accumulation could potentially prevent or delay Parkinson’s disease onset and disease progression.

DA oxidation to [o-]quinones is an essential event required for NM synthesis^6–9^. In particular, NM is formed by oxidation of excess cytosolic DA that is not accumulated into synaptic vesicles by the vesicular monoamine transporter-2 (VMAT2; SLC18A2), an H^+^- ATPase antiporter found on the membrane of secretory vesicles of presynaptic neurons that rapidly internalizes into synaptic vesicles both recently-synthesized DA and extracellular DA reuptaken by the dopamine transporter (DAT)^10–12^. VMAT2-mediated DA vesicular uptake is essential to prepare DA for subsequent release and to prevent its cytosolic accumulation and oxidation into potentially toxic [o-]quinones, the latter being rapidly transformed into eumelanic or pheomelanic intermediates that constitute the melanic component of NM^6–9^. In line with this, there is an inverse relationship between NM content and VMAT2 immunoreactivity in human midbrain DA neurons^13^, with neurons exhibiting the highest VMAT2 levels being the ones with the lowest NM levels, and the least vulnerable to PD-linked neurodegeneration. Conversely, the most vulnerable ventral SNpc neurons accumulate the highest NM levels and exhibit the lowest VMAT2 levels^13^. Supporting a potential pathogenic perturbation of VMAT2 in Parkinson’s disease: (i) postmortem Parkinson’s disease brains exhibit decreased VMAT2 expression in nigrostriatal DA terminals^14^, (ii) VMAT2-dependent DA encapsulation within synaptic vesicles is defective in early Parkinson’s disease cases^15^, (iii) VMAT2 mRNA is decreased in circulating platelets from Parkinson’s disease patients, suggesting a systemic VMAT2 deficiency^16^, (iv) gain-of-function VMAT2 variants in humans have been associated to decreased likelihood of developing Parkinson’s disease^17,18^. By taking advantage of our newly developed NM-producing animal model, here we assessed whether enhancement of DA vesicular encapsulation by VMAT2 overexpression may provide therapeutic benefit in the context of Parkinson’s disease by maintaining DA homeostasis, decreasing potentially toxic DA oxidized species and preventing age-dependent NM accumulation above its pathogenic threshold.

## Materials and methods

### Study Design

In this study we aimed to determine whether VMAT2 overexpression could decrease age-related NM production *in vivo* and prevent/attenuate Parkinson’s disease pathology. This question was addressed by taking advantage of the only currently available rodent model of NM production, which we have recently developed, based on the unilateral viral vector-mediated overexpression of TYR in the rat SNpc^5^. In parallel to NM accumulation, these animals develop a progressive Parkinson’s disease-like phenotype characterized by motor deficits, LB-like inclusion formation, nigrostriatal neurodegeneration, extracellular NM released from dying neurons and neuroinflammation. Adult male Sprague–Dawley rats were randomly distributed into three different experimental groups receiving, respectively, unilateral intranigral injections of either AAV-TYR, AAV-VMAT2 or a combination of both. After confirming that the combination of both vectors did not interfere with the expression of one another, we evaluated in all groups behavioral (motor asymmetry), histological (intracellular NM levels, LB-like inclusion formation, TH & VMAT2 downregulation, nigrostriatal degeneration, extracellular NM, neuroinflammation) and metabolic (DA vesicular uptake, DA levels, DA metabolism, DA oxidation) changes. Based on our previous observations, two different experimental time-points were selected for these evaluations: (i) at the onset of NM-linked neuronal dysfunction but before neurodegeneration (i.e. 2m post-AAV), equivalent to prodromal Parkinson’s disease; (ii) once nigrostriatal neurodegeneration is fully established in these animals (i.e. 6m post-AAV), equivalent to established Parkinson’s disease^5^. Two types of controls were used for the different quantifications: (1) the unaffected contralateral (non-AAV injected) hemisphere of each animal, representing an internal control/baseline; (2) a group of rats unilaterally injected with AAV-VMAT2 in the SN, which serve as a control of the potential effects of VMAT2 overexpression by itself. In addition, we had previously reported that unilateral nigral injections of the corresponding AAV-empty vector (EV) or vehicle/sham does not produce any nigral pathology in these animals in contrast to AAV-TYR injections^5^. Therefore, AAV-EV or vehicle/sham injections were not used in this study, to minimize the number of animals according to international regulations on the protection of animals used for experimental purposes. All researchers were blinded to the experimental groups analyzed.

### Animals

Adult male Sprague–Dawley rats (Charles River), 225–250 g at the time of surgery, were housed two to three per cage with ad libitum access to food and water during a 12 hours (h) light/dark cycle. All the experimental and surgical procedures were conducted in accordance with the European (Directive 2010/63/UE) and Spanish laws and regulations (Real Decreto 53/2013; Generalitat de Catalunya Decret 214/97) on the protection of animals used for experimental and other scientific purposes, and approved by the Vall d’Hebron Research Institute (VHIR) Ethical Experimentation Committee, to ensure the use of the minimum necessary number of animals and the application of protocols that cause the least pain, suffering or distress to animals. Rats were randomly distributed into the different experimental groups and control and experimental groups were processed at once to minimize bias.

### Stereotaxic infusion of viral vectors

Recombinant AAV vector serotype 2/1 expressing the human tyrosinase cDNA driven by the cytomegalovirus (CMV) promoter (AAV-TYR; concentration 2.6 × 10^13^ gc/mL) and AAV serotype 2/9 containing the human VMAT2 cDNA fused to 3 Flag epitopes under control of the CMV promoter (AAV-VMAT2; concentration 7.2 × 10^12^ gc/mL) were produced at the Viral Vector Production Unit of the Autonomous University of Barcelona (UPV-UAB, Spain). Surgical procedures were performed with the animals placed under general anesthesia using isoflurane (5% for the induction phase and 2% for the maintenance phase) (Baxter). Vector solutions were injected using a 10 μL Hamilton syringe fitted with a glass capillary (Hamilton model Cat#701). Animals received 2 μL of a 0.65:1.35 mixture of AAV-TYR+AAV-Flag-VMAT2, AAV-TYR+vehicle or vehicle+AAV-Flag-VMAT2 to achieve a final titration of 1.7 × 10^13^ gc/mL for AAV-TYR and 9.7 × 10^12^ gc/mL for AAV-VMAT2. In the TYR/vehicle and VMAT2/vehicle groups, viral vectors were diluted with vehicle (PBS-MK/40% Iodixanol). Infusion was performed at a rate of 0.4 μL/min and the needle was left in place for an additional 4 min period before it was slowly retracted. Injection was carried out unilaterally on the right side of the brain at the following coordinates (flat skull position), right above the SNpc: antero-posterior: -5.29 mm; medio-lateral: -2 mm; dorso-ventral: -7.6 mm below dural surface, calculated relative to bregma according to the stereotaxic atlas of Paxinos and Watson.

### Brain processing for UPLC-MS/MS and histological analyses

Two or six months after injection of viral vectors animals were euthanized and the brains quickly removed and placed over a cold plate. For histological analyses, the posterior portion of the brain was embedded in ice-cold formaldehyde solution 4% phosphate buffered (Panreac) for 24 h and subsequently processed for paraffin embedding following standard procedures. Sectioning was performed with a sliding microtome (Leica, Germany) at 5 µm-thickness. For UPLC-MS/MS analyses, contralateral and ipsilateral striata (Str) were dissected, frozen on dry ice, and stored at -80°C until use. In an additional set of animals euthanized at two months, Str and ventral midbrain (vMB) were dissected, frozen on dry ice, and stored at -80°C until analyzed by UPLC-MS/MS. Two additional sets of animals were euthanized at one month post-AAV injection and their brains processed for either histology or qPCR, respectively, to assess transduction efficiency of the viral vectors, alone or in combination.

### Immunohistochemistry

Deparaffinized rat brain sections were quenched for 10 min in 3% H_2_O_2_-10% (vol/vol) methanol. Antigen retrieval in paraffin sections was performed with a 10 mM citric acid solution at pH 6.0 in a microwave for 20 min. Sections were rinsed 3 times in 0.1 M Tris buffered saline (TBS) between each incubation period. Blocking for 1 h with 5% (vol/vol) normal goat serum (NGS, Vector Laboratories) was followed by incubation with the appropriate primary antibody at 4°C for 48 h in 2% (vol/vol) serum. Details about primary antibodies used for immunohistochemistry can be found in Table 1. Sections were then incubated with the corresponding secondary biotinylated antibody (Vector Laboratories), visualized by incubation with avidin-biotin-peroxidase complex (Immunopure ABC Peroxidase staining kit; Thermo Fisher Scientific), using the VectorSG Peroxidase Substrate Kit (Vector Laboratories) as a chromogen, and mounted and coverslipped with DPX mounting medium (Sigma-Aldrich).

### Immunofluorescence

Immunofluorescence procedure was a similar to the previously reported immunohistochemistry protocol without the quenching step. Blocking was performed with 5% (vol/vol) NGS and 0.1% (vol/vol) Triton X-100 (Sigma-Aldrich) in phosphate buffered saline (PBS) solution. Corresponding primary antibodies were incubated together overnight at 4°C in 2% (vol/vol) serum. Details about primary antibodies used for immunofluorescence can be found in Table 1. Adequate Alexa 488, 569, and/or 647-conjugated secondary antibodies (1:1000, Thermo Fisher Scientific) were incubated simultaneously for 1 h at RT in 2% (vol/vol) serum. Nuclei were stained with Hoechst 33342 (1:2000, Thermo Fisher Scientific) in 1x PBS for 10 min. Sections were coverslipped using the Dako Cytomation Fluorescent Mounting Medium (Dako).

Immunofluorescent images were taken with a LSM 980 with Airyscan 2 confocal microscope (Zeiss, Germany) and were analyzed with ZEN 3.1 software (Zeiss, Germany). The total number of p62-positive cytoplasmic inclusions was manually determined in one SNpc section per animal, from two different experimental groups: TYR (*n*=8) and TYR+VMAT2 (*n*=7), at two months post-AAV injections. All quantifications were performed by an investigator blinded to the experimental groups.

### Cell counting

Assessment of the total number of TH-positive neurons, the number of NM-laden neurons (with or without TH) and extracellular NM aggregates in the SNpc was performed by stereology according to the fractionator principle, using the MBF Bioscience StereoInvestigator 11 (64 bits) Software (Micro Brightfield) coupled to a Zeiss Imager.D1 microscope with an AxioCam MRc camera (Zeiss, Germany). Serial 5 µm-thick paraffin sections covering the entire SNpc were included in the counting procedure (every 17^th^ section for a total of 10-12 sections analyzed/animal, Supplementary Figure 1). The following sampling parameters were used: (i) a fixed counting frame with a width and length of 50 μm; (ii) a sampling grid size of 115 x 70 μm and (iii) a multiplication factor of 17. The counting frames were placed randomly by the software at the intersections of the grid within the outlined structure of interest. Objects in both brain hemispheres were independently counted following the unbiased sampling rule using a 100x lens and included in the measurement when they came into focus within the dissector. A coefficient of error of <0.10 was accepted. Data for the total numbers of TH-positive neurons and NM-containing neurons are expressed as absolute numbers for each hemisphere. The total number of SNpc DA neurons was calculated by considering all TH^+^NM^+^, TH^-^NM^+^ and TH^+^NM^-^ neurons. The percentage of TH downregulation was calculated by considering the total number of TH^+^NM^+^ and the total number of TH^-^NM^+^ with respect to the total number of neurons containing NM. All quantifications were performed by an investigator blinded to the experimental groups. For the assessment of the number of VMAT2-positive neurons, a similar approach was applied to a selected SNpc middle section/animal. The percentage of ipsilateral VMAT2 downregulation was calculated by considering the number of VMAT2^-^NM^+^ neurons vs total SNpc DA neurons (including VMAT2^+^NM^+^, VMAT2^+^NM^-^ and VMAT2^-^NM^+^ neurons).

For the quantification of VMAT2-Flag and TYR transduction efficiency, high resolution micrographs were acquired with an Olympus Slideview VS200 slide scanner and the Olyvia 3.3 software (Olympus, Japan). A specific artificial intelligence (AI)-assisted algorithm was designed for the quantification of vector SG immunostained neurons (VMAT2-Flag, TYR or TH, Supplementary Figure 2) using the Olympus V200 Desktop 3.3 software. Adjacent serial 5 µm-thick paraffin sections covering the entire SNpc were included in the counting procedure (every 17^th^ section for a total of 10-12 sections analyzed/animal, Supplementary Figure 1). Each automatized counting was manually revised by an investigator blinded to the experimental groups.

### Quantification of neuroinflammation parameters

Quantification of Iba-1, CD68 and GFAP-positive cells was performed in SNpc sections adjacent to those used for stereological cell counts. Serial 5 µm-thick paraffin sections covering the entire SNpc (every 17^th^ section for a total of 10-12 sections analyzed/animal, Supplementary Figure 1) were scanned using the Panoramic Midi II FL, HQ SCIENTIFIC 60x scanner and section images were acquired with CaseViewer software (3D Histech, Hungary). For quantification of Iba-1, CD68 and GFAP-positive cells, specific AI-based algorithms were created using the Aiforia platform (Aiforia Technologies, Finland). Iba-1-positive cells were counted separately in 2 different groups according to their activation state: non-reactive (branched) and reactive (amoeboid). CD68 and GFAP-positive cells were counted individually. Data is presented as the number of positive cells per quantified area (in mm^2^). All quantifications were performed by an investigator blinded to the experimental groups. For illustration purposes, high resolution micrographs were acquired with the Olympus Slideview VS200 slide scanner and the Olyvia 3.3 software.

### Intracellular NM quantification

Intracellular NM levels were quantified in both AAV-TYR and AAV-TYR+AAV-VMAT2-injected animals at two months post-AAV injections in 5 µm-thick paraffin-embedded H&E-stained sections covering the whole SNpc for each animal. In these sections, SNpc dopaminergic neurons were identified by the visualization of unstained NM brown pigment. Midbrain sections were scanned using the Pannoramic Midi II FL, HQ SCIENTIFIC 60x and section images were acquired with CaseViewer software at an objective magnification of 63x. For illustration purposes, high resolution micrographs were acquired with an Olympus Slideview VS200 slide scanner and the Olyvia 3.3 software. All NM-positive neurons in a representative SNpc section of each animal were analyzed by means of optical densitometry using ImageJ software (NIH, USA) to quantify the intracellular density of NM pigment, as previously reported^5^. The pixel brightness values for all individual NM-positive cells (excluding the nucleus) in all acquired images were measured and corrected for non-specific background staining by subtracting values obtained from the neuropil in the same images. All quantifications were performed by an investigator blinded to the experimental groups.

### Chromatographic determination of dopaminergic metabolites in brain samples

UPLC-MS/MS analysis of rat brain regions was performed using our previously validated method^19^ with some modifications. Each sample was injected three times into the UPLC-MS/MS system to analyze different sets of compounds i.e. MIX1, MIX2 and MIX3. MIX1 includes dopamine (DA), 3-methoxytyramine (3MT), 3,4-dihydroxyphenylalanine (L-DOPA) and aminochrome (AC); MIX2 includes 3,4-Dihydroxyphenylacetic acid (DOPAC) and MIX3 includes 5-*S*-cysteinyldopa (5SCD) and 5-*S*-cysteinyldopamine (5SCDA). IS was added to every set of compounds. 5SCD and 5SCDA standards were kindly donated by Professor Kazumasa Wakamatsu and Professor Shosuke Ito at the Fujita Health University, Aichi, Japan. Aminochrome standard (0.5 mM) was freshly prepared as previously described ^19,20^. Multiple Reaction Monitoring (MRM) acquisition settings for the targeted metabolites are summarized in Table 2. Samples with a concentration between LOD and LOQ or bigger than LOQ were considered acceptable; samples with a concentration lower than LOD were considered as the LOD value. Data was normalized by the protein concentration and presented as the percentage of the contralateral concentration or ratio.

### Cylinder behavioral test

Rats were tested for left and right forepaw use with the cylinder test one week before surgery and at two and six months after viral injection. The number of animals for each condition was VMAT2 (*n*=7), TYR (*n*=8) and TYR+VMAT2 (*n*=7-8). For the performance of the cylinder test, rats were first allowed to habituate to the experimental room for at least 1 h before each test. Then, rats were put in a glass cylinder and the total number of left and right forepaw touches performed within 5 min was counted. Data are presented as the percentage of the contralateral paw usage. Behavioral equipment was cleaned with 70% ethanol after each test session to avoid olfactory cues. All behavioral tests were performed during the light cycle by an investigator blinded to the experimental groups.

### Human tyrosinase gene expression

Total RNA was extracted from dissected ipsilateral and contralateral vMB from AAV-TYR- injected rats using mirVana PARIS RNA and Native Protein Purification Kit (Thermo Fisher Scientific # AM1556). RNA concentration was determined using a NanoDrop ND-1000 Spectrophotometer. 0.5 µg of total RNA were retrotranscribed using High-Capacity cDNA Reverse Transcription Kit (Thermo Fisher Scientific # 4368814). qPCR was performed with 10 ng of cDNA per well in technical triplicates mixed with Taqman Gene Expression Master Mix (Applied Biosystems, # 4369016) and Taqman gene expression assays [human tyrosinase (TYR) (Hs00165976_m1, Applied Biosystems) using standard procedures in a 7900HT Fast Real Time Instrument (Applied Biosystems). Thresholds cycles (Cts) for each target gene were normalized to an endogenous reference gene (Gapdh Mm99999915_g1, Rpl19 Mm02601633_g1 and Ppia Mm02342430_g1). Water was included in the reaction as a non-template (negative) control. The relative expression was calculated with the ΔCt-method.

### Statistical analysis

Statistical analyses were performed with GraphPad Prism software (v6, GraphPad Software Inc, USA). No statistical methods were used to pre-determine sample size but our sample sizes are equivalent to those reported in previous similar publications^5^. Outlier values were identified by the ROUT test and excluded from the analyses when applicable. Selection of the pertinent data representation and statistical test for each experiment was determined after formally testing for normality with the Shapiro-Wilk normality test. Accordingly, differences among means or medians were analyzed either by 1- or 2-way analysis of variance (ANOVA), Kruskal–Wallis ANOVA on ranks or Mann–Whitney rank sum test, as appropriate and indicated in each figure legend. When ANOVA showed significant differences, pairwise comparisons between means were subjected to Tukey’s post-hoc testing for multiple comparisons. Values are expressed either as mean ± standard error of the mean (SEM) or presented as box plots, with minimum, maximum and median indicated, depending on the performance of parametric or non-parametric analyses, respectively. In all analyses, the null hypothesis was rejected at the 0.05 level.

### Data availability

All data are available in the main text or the supplementary materials.

## Results

### VMAT2 overexpression reduces catechol oxidation, NM production and LB-like inclusion formation in humanized rats

To determine first whether VMAT2 overexpression was able to boost DA vesicular uptake *in vivo*, adult rats received a single unilateral stereotaxic injection of an adeno-associated viral vector (AAV) expressing flagged human VMAT2 (AAV-VMAT2) above the right SNpc (Fig. 1A). In these animals, conspicuous exogenous VMAT2 expression was observed in ipsilateral SNpc and striatum at 1 month (m) post-AAV injection, as assessed by Flag immunohistochemistry (Fig. 1B). Overexpressed VMAT2 proved to be functionally active, as demonstrated by (i) an enhanced striatal DA storage accompanied by a decreased DA metabolism^21^ and (ii) a ∼2-fold increase in striatal DA:L-DOPA ratio, an index of DA vesicular uptake^22^, measured by ultra-performance liquid chromatography-tandem mass-spectrometry (UPLC-MS/MS) at 2m post-AAV injection (Fig. 1C and Supplementary Table 1).

**Figure 1.**
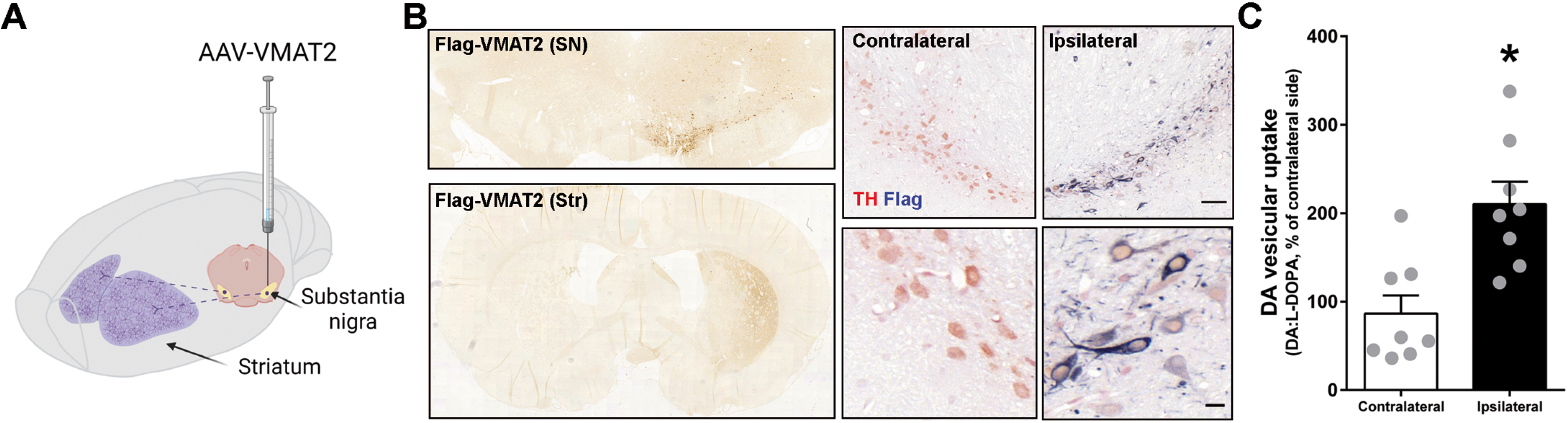
Overexpression of functional VMAT2 in the nigrostriatal pathway of rats. (**A**) Schematic representation of the site of AAV-VMAT2 unilateral stereotaxic injection above the SN of the rat brain. (**B**) L*eft,* representative images of nigral (top) and striatal (bottom) Flag-VMAT2 immunolabeling ipsilateral to the injection in AAV-VMAT2-injected rats at 1 m post-AAV injection. *Right,* representative images of TH-Flag colocalization in ipsilateral SNpc of AAV-VMAT2-injected rats at 1 m post-AAV injection; Scale bar: 50 µm (top), 10 µm (bottom). (**C**) Quantification of striatal DA vesicular uptake by UPLC-MS/MS in AAV-VMAT2-injected rats at 2m post-AAV injection, as calculated by the ratio between striatal DA:L-DOPA levels^22^. Values are mean ± SEM. \**p<*0.05 vs. contralateral (non-injected) side; Two-tailed t-test. *N=*8 animals per group. Drawing in A was created with BioRender.com.

After confirming that we are able to overexpress functional VMAT2 *in vivo*, we next assessed whether nigrostriatal VMAT2 overexpression in NM-producing rats could increase DA vesicular uptake and reduce the rate of non-encapsulated DA oxidation into NM precursors. To address this question, AAV-VMAT2 and AAV-TYR were co-injected above the rat SNpc (Fig. 2A, Supplementary Figure 2, Supplementary Table 2), with additional groups of animals receiving equivalent amounts of either AAV-TYR or AAV-VMAT2 separately. In AAV-TYR-injected rats, we have previously reported that intracellular NM starts reaching pathological levels at 2m post-AAV injection, at which time these animals exhibit early functional defects, including impaired DA release and motor deficits, prior to degeneration^5^. Here we found, similar to Parkinson’s disease patients^15,23^, that early functional changes in AAV-TYR melanized rats are accompanied by a downregulation of endogenous vmat2 levels, as shown by a reduction of vmat2 expression within NM-containing neurons (Fig. 2A and Supplementary Table 3), and alterations in nigrostriatal DA homeostasis, such as reduced DA vesicular uptake, decreased DA levels, increased DA metabolism and enhanced catechol oxidation (Fig. 2B-E, Supplementary Figure 3 and Supplementary Tables 4&5). Remarkably, all these pathological changes were prevented by overexpressing VMAT2 in these animals (Fig. 2A-E, and Supplementary Tables 4&5).

**Figure 2.**
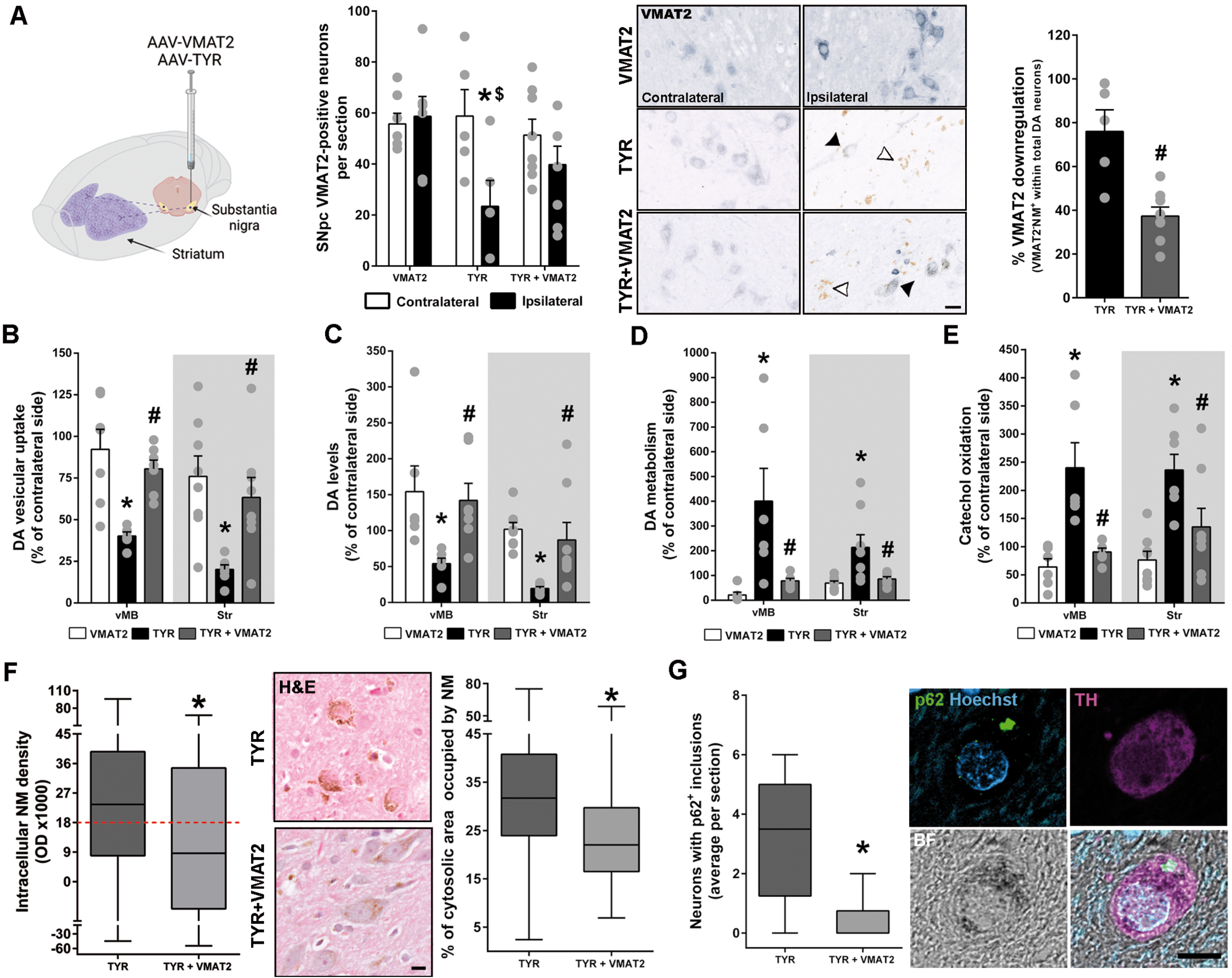
Reduced catechol oxidation, NM accumulation and LB-like pathology by VMAT2 overexpression in NM-producing rats. (**A**) *Left*, schematic representation of the site of AAV-VMAT2, AAV-TYR or AAV-VMAT2+AAV-TYR unilateral stereotaxic injection above the SN of the rat brain. *Center-left*, quantification of contralateral and ipsilateral SNpc neurons immunopositive for VMAT2 (including VMAT2^+^NM^+^ and VMAT2^+^NM^-^) in AAV-VMAT2, AAV-TYR and AAV-TYR+AAV-VMAT2-injected rats at 2m post-AAV injections. *Center-right*, representative contralateral and ipsilateral VMAT2-immunostained SNpc sections from these animals are shown in micrographs; Scale bar 20 µm *Right*, ipsilateral VMAT2^-^NM^+^ neurons vs total DA neurons (including VMAT2^+^NM^+^, VMAT2^+^NM^-^ and VMAT2^-^NM^+^ neurons) as an index of VMAT2 downregulation. (**B-E**) UPLC-MS/MS quantification of: (**B**) DA vesicular uptake (calculated as DA:L-DOPA); (**C**) DA levels; (**D**) DA metabolism (calculated as [3MT+DOPAC]:DA); (**E**) catechol oxidation (calculated as [5SCD:L-DOPA]+[(5SCDA+AC):DA]) in ventral midbrain (vMB) and striatal samples from AAV-VMAT2, AAV-TYR and AAV-TYR+AAV-VMAT2-injected animals at 2 m post-AAV injections. Results represent ipsilateral levels shown as the percentage of the contralateral (non-injected) side. Values are mean ± SEM. (**F**) *Left*, quantification of intracellular NM levels by optical densitometry in ipsilateral SNpc DA neurons of AAV-TYR and AAV-TYR+AAV- VMAT2-injected rats at 2m post-AAV injections. Dotted red line indicates the previously reported pathogenic threshold of intracellular NM accumulation^5^. *Center*, representative images of 5 µm-thick H&E-stained nigral sections from these animals (unstained NM in brown); Scale bar: 10 µm. *Right*, percentage of the neuronal cytosolic area occupied by NM in AAV-TYR and AAV-TYR+AAV-VMAT2-injected rats at 2 m post-AAV injections. Values are median + min to max. (**G**) *Left*, quantification of the number of NM-filled neurons exhibiting p62-positive LB-like cytosolic inclusions in AAV-TYR and AAV-TYR+AAV- VMAT2-injected rats at 2m post-AAV injections. Values are median + min to max. *Right*, representative confocal images of a melanized TH^+^ neuron with a LB-like inclusion immunopositive for p62 in the SNpc of a AAV-TYR-injected rat; BF, brightfield; Scale bar: 10 µm. In **A**, *center-left* \**p*<0.05 vs. respective contralateral side; ^$^*p<*0.05 vs. ipsilateral VMAT2; Two-way ANOVA with Tukeýs post-hoc test; *center-right* ^#^*p<*0.05 vs. TYR; Two-tailed t-test *N=* 7 (VMAT2), 5 (TYR), 8 (TYR+VMAT2) animals. In **B-E**, \**p<*0.05 vs. VMAT2; ^#^*p<*0.05 vs. TYR; one-way ANOVA with Tukeýs post-hoc test. N= 6-8 (VMAT2), 6-8 (TYR), 6-8 (TYR+VMAT2) animals. In **F-G**, \**p*<0.05 vs. TYR; Mann-Whitney test. *N=*594 neurons from 8 AAV-TYR-injected animals and 180 neurons from 8 AAV- TYR+AAV-VMAT2-injected animals. Drawing in A was created with BioRender.com.

When oxidized, tyrosine, L-DOPA or DA produce dopaquinone or dopamine-o-quinone that will act as precursors for either the eumelanin (brown) and/or the pheomelanin (reddish) melanic components of NM (Supplementary Figure 3). Thus, we next assessed whether attenuated catechol oxidation in VMAT2/TYR co-injected animals was associated to a reduced NM production. Quantification of intracellular NM optical density within individual SNpc neurons, which reflects NM levels^3,24^, revealed that intracellular NM density was significantly decreased in SNpc neurons of TYR+VMAT2 rats, compared to AAV-TYR-injected animals (Fig. 2F). While intracellular NM optical density could be influenced by potential changes in NM composition, independently of the actual levels of NM, we also observed a significant reduction in the percentage of cytosolic neuronal area occupied by NM in TYR+VMAT2 animals compared to their TYR counterparts (Fig. 2F), indicating an actual decrease of NM levels in these animals. Importantly, VMAT2 overexpression reduced intracellular NM to levels below the pathogenic NM threshold previously identified in NM-producing animals^5^ (red dotted line in Fig. 2F). Above this threshold, in addition to the functional defects reported earlier, NM-laden neurons develop LB-like inclusions (Fig. 2G and Supplementary Table 3), a hallmark of Parkinson’s disease pathology closely associated to NM accumulation^5,25,26^. LB-like inclusions in NM-producing rats are immunopositive for p62, a common component of LBs in Parkinson’s disease brains^27^, and were previously reported to peak at 2m post-AAV-TYR injection, thus coinciding with functional alterations and preceding neurodegeneration in these animals^5^. Importantly, the reduction of intracellular NM levels by VMAT2 overexpression was accompanied with attenuated LB-like inclusion formation in these neurons (Fig. 2G and Supplementary Table 3). Taken together, these results demonstrate that VMAT2 overexpression is able to restore DA homeostasis, reduce NM production/accumulation and attenuate LB-like pathology *in vivo*.

### VMAT2 overexpression prevents NM-linked neurodegeneration

In NM-producing rats, early functional defects observed at 2m are followed by progressive nigrostriatal neurodegeneration beginning at 4m post-AAV-TYR injection^5^. To determine whether decreased DA oxidation and subsequent reduction in NM accumulation achieved by VMAT2 overexpression were associated with preserved neuronal integrity, we assessed the long-term effects of VMAT2 overexpression in NM-producing rats at 6m post-AAV-TYR injection, once nigrostriatal degeneration is well-established in this animal model^5^. By this time, NM-producing rats exhibited a ∼90% reduction in ipsilateral SNpc TH-positive neurons that was markedly attenuated by concomitant VMAT2 overexpression (Fig. 3A and Supplementary Table 6). Similar to Parkinson’s disease brains, NM-producing rats also exhibited a phenotypic loss of TH expression within NM-laden neurons, which reflects neuronal dysfunction at early stages of neurodegeneration^1^, as shown by an increased percentage of TH-immunonegative neurons within the total population of NM-containing neurons (Fig. 3B and Supplementary Table 6). Interestingly, VMAT2 overexpression greatly attenuated TH downregulation occurring within pigmented neurons, indicating a functional preservation of these neurons (Fig. 3B and Supplementary Table 6). To distinguish between the effects of VMAT2 on TH downregulation and on actual cell death, we assessed the number of total SNpc DA neurons (including TH-immunopositive and TH-immunonegative pigmented SNpc neurons), which confirmed the ability of VMAT2 overexpression to prevent NM-linked SNpc DA neurodegeneration in NM-producing rats (Fig. 3C and Supplementary Table 6). Overall, these results indicate that VMAT2 overexpression is able to prevent neuronal dysfunction and degeneration linked to excessive NM production/accumulation.

**Figure 3.**
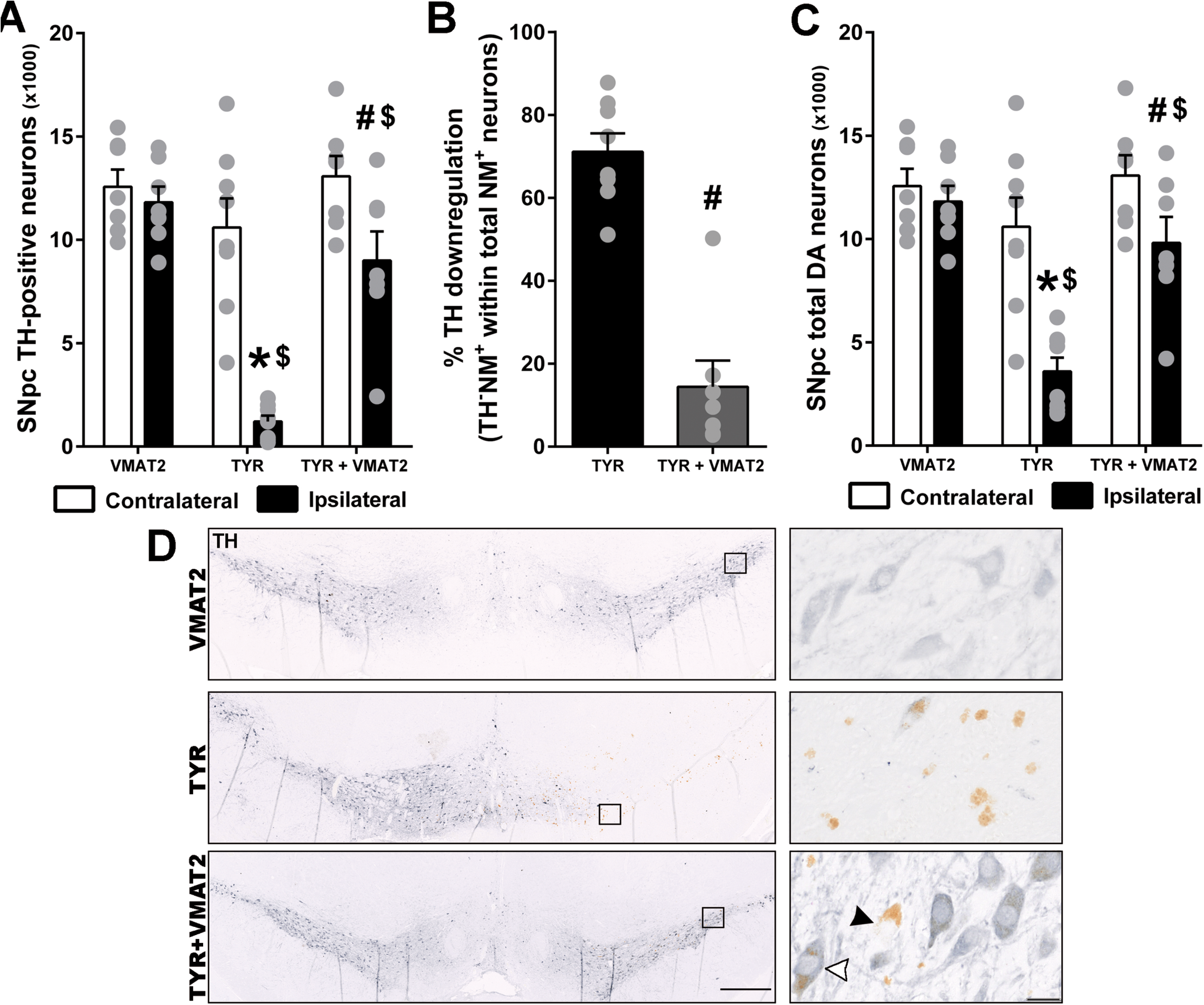
VMAT2 overexpression prevents SNpc neurodegeneration in NM-producing rats. Stereological nigral cell counts of: (**A**) TH^+^ neurons; **(B)** TH^-^NM^+^ neurons vs total NM^+^ neurons (including TH^+^NM^+^ and TH^-^NM^+^ neurons) as an index of TH downregulation; (**C**) total DA neurons (including TH^+^NM^+^, TH^+^NM^-^ and TH^-^NM^+^ neurons), in AAV-VMAT2, AAV-TYR and AAV-TYR+AAV-VMAT2-injected rats at 6m post-AAV injections. (**D**) Representative images of TH-positive neurons (TH in blue, unstained NM in brown), in which TH^-^NM^+^ (black arrowhead) and TH^+^NM^+^ (white arrowhead) neurons are indicated for identification purposes; scale bars: 200 µm & 20 µm (inset). Values are mean ± SEM. \**p*<0.05 vs. ipsilateral VMAT2; ^#^*p<*0.05 vs. ipsilateral TYR; ^$^*p<*0.05 vs. respective contralateral side; ^&^p<0.05 vs. TH^+^NM^+^; Two-way ANOVA with Tukeýs post-hoc test. *N=*7 (VMAT2), 8 (TYR) and 7 (TYR+VMAT2) animals.

### Attenuated neurodegeneration by VMAT2 overexpression is associated to decreased extracellular NM debris and inflammation

Concomitant with SNpc pigmented cell death, both PD brains and NM-producing rats exhibit abundant extracellular NM released from dying neurons, some of which is surrounded by, or contained within activated microglia, indicative of an active, ongoing neurodegenerative process (i.e. neuronophagia)^5,28^. In this context, the marked neuroprotective effect provided by VMAT2 overexpression in NM-producing rats at 6m post-AAV injection was associated to a drastic reduction of extracellular NM in these animals (Fig. 4A). Microglial cells are the main players in the recognition, engulfment and clearance of extracellular NM^28,29^. In agreement with this, the number of microglial cells with non-reactive (ramified) and phagocytic/reactive (large amoeboid de-ramified) morphology was markedly increased in AAV-TYR-injected rats in association to extracellular NM and concomitant to cell death (Fig. 4B and Supplementary Table 6). By attenuating neurodegeneration and subsequent accumulation of extracellular NM, VMAT2 overexpression was associated to a diminished microglial activation in these animals (Fig. 4B and Supplementary Table 6). Similarly, VMAT2 overexpression also precluded the recruitment of CD68-positive cells occurring in NM-producing animals, corresponding to tissue-resident or blood-borne macrophages with phagocytic activity that are found in close association with extracellular NM (Fig. 4C and Supplementary Table 6). Additional inflammatory changes observed in NM-producing rats included increased astrocyte reactivity, widely distributed within the SNpc, which was also markedly attenuated by VMAT2 overexpression (Fig. 4D). Taken together, these results reveal that, by reducing cell death and subsequent NM release from dying neurons, VMAT2 overexpression is able to prevent the overall PD-like inflammatory response linked to the accumulation of extracellular NM.

**Figure 4.**
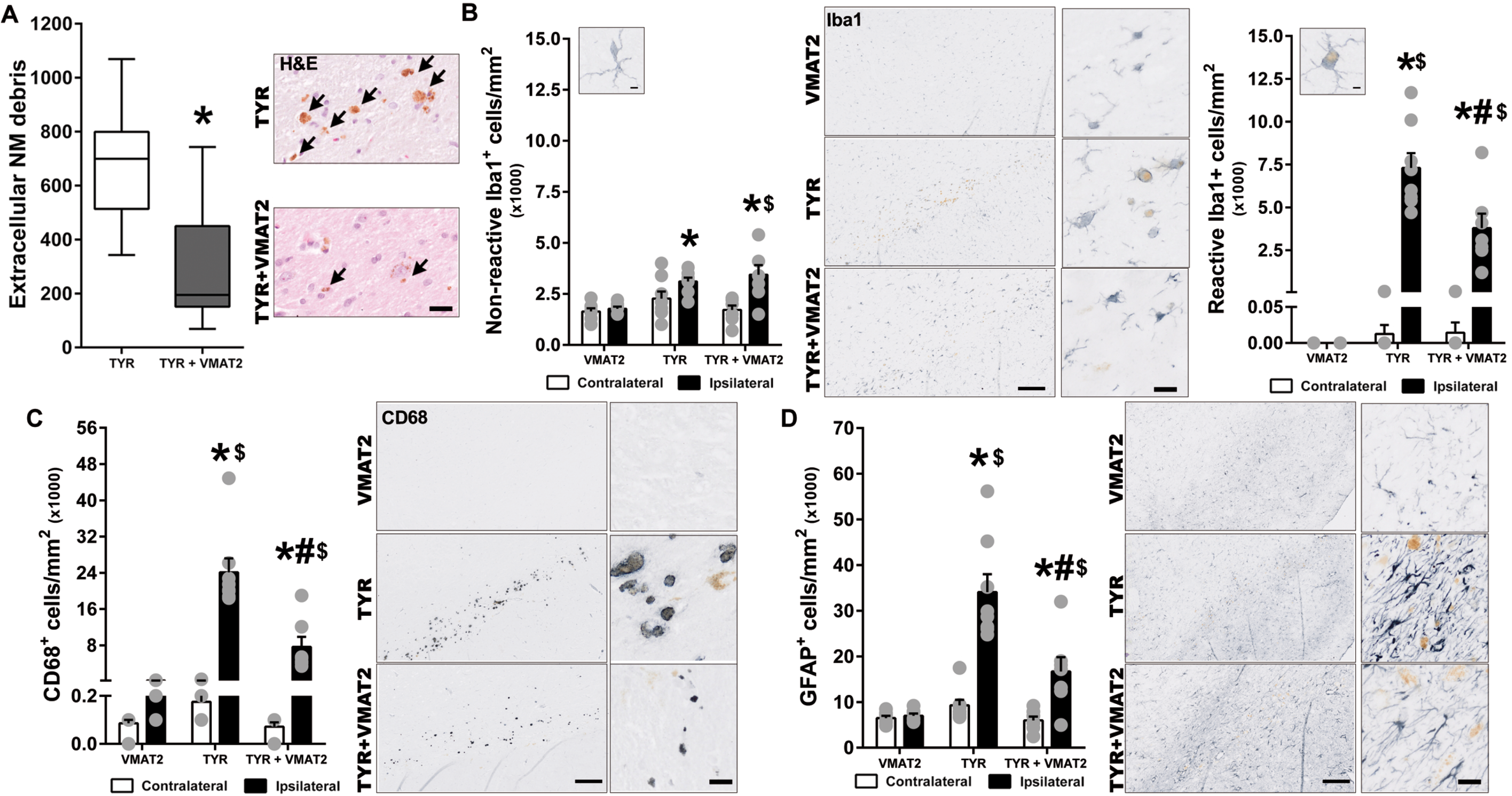
VMAT2 overexpression reduces extracellular NM debris and inflammation in the SNpc of NM-producing rats. Quantification of: (**A**) extracellular NM clusters; (**B**) Iba1+ non-reactive (left) and reactive (right) microglia; (**C**) CD68+ cells; (**D**) GFAP+ astrocytes, in AAV-VMAT2, AAV-TYR and AAV-TYR+AAV-VMAT2-injected rats at 6m post-AAV injections. Photomicrographs correspond to representative images of (**A**) H&E-stained SNpc sections (unstained NM in brown) and (**B-D**) immunostained SNpc sections for Iba1, CD68 or GFAP (in blue, unstained NM in brown). Scale bars: 10 µm (**A**, insets in **B-D**), 200 µm (B-D). In (**A**) Values are median + min to max. In **B-D**, values are mean ± SEM. \**p*<0.05 vs. ipsilateral VMAT2; ^#^p<0.05 vs. ipsilateral TYR; ^$^p<0.05 vs. contralateral side; Mann-Whitney test **(A)** or two-way ANOVA with Tukeýs post-hoc test (**B-D**). *N=* 7 (VMAT2), 8 (TYR), 7 (TYR+VMAT2) animals.

### Preservation of striatal DA levels and metabolism by VMAT2 overexpression prevents motor deficits in NM-producing rats

We next determined whether the neuroprotective effects of VMAT2 overexpression on SNpc DA neuron cell bodies were accompanied by a preservation of striatal DA levels in NM-producing rats. At 6m post-AAV-TYR injections, concomitant to SNpc neurodegeneration, NM-producing rats exhibited a marked reduction of striatal DA levels as assessed by UPLC-MS/MS (Fig. 5A and Supplementary Table 7). Consistent with its effect at preserving DA nigral cell bodies, VMAT2 overexpression also prevented the loss of striatal DA in these animals (Fig. 5A and Supplementary Table 7). The preservation of striatal DA levels by VMAT2 overexpression was associated to a re-established striatal DA vesicular uptake, reduced DA metabolism and decreased catechol oxidation in these animals (Fig. 5B-D, and Supplementary Table 7). Confirming a functional preservation of the nigrostriatal circuit, VMAT2 overexpression prevented contralateral forepaw hypokinesia occurring in NM-producing rats, both at 2m and 6m post-AAV injections (Fig. 5E). Collectively, these results demonstrate the ability of VMAT2 overexpression to provide a morphological and functional long-term preservation of the nigrostriatal pathway *in vivo* in a humanized Parkinson’s disease model.

**Figure 5.**
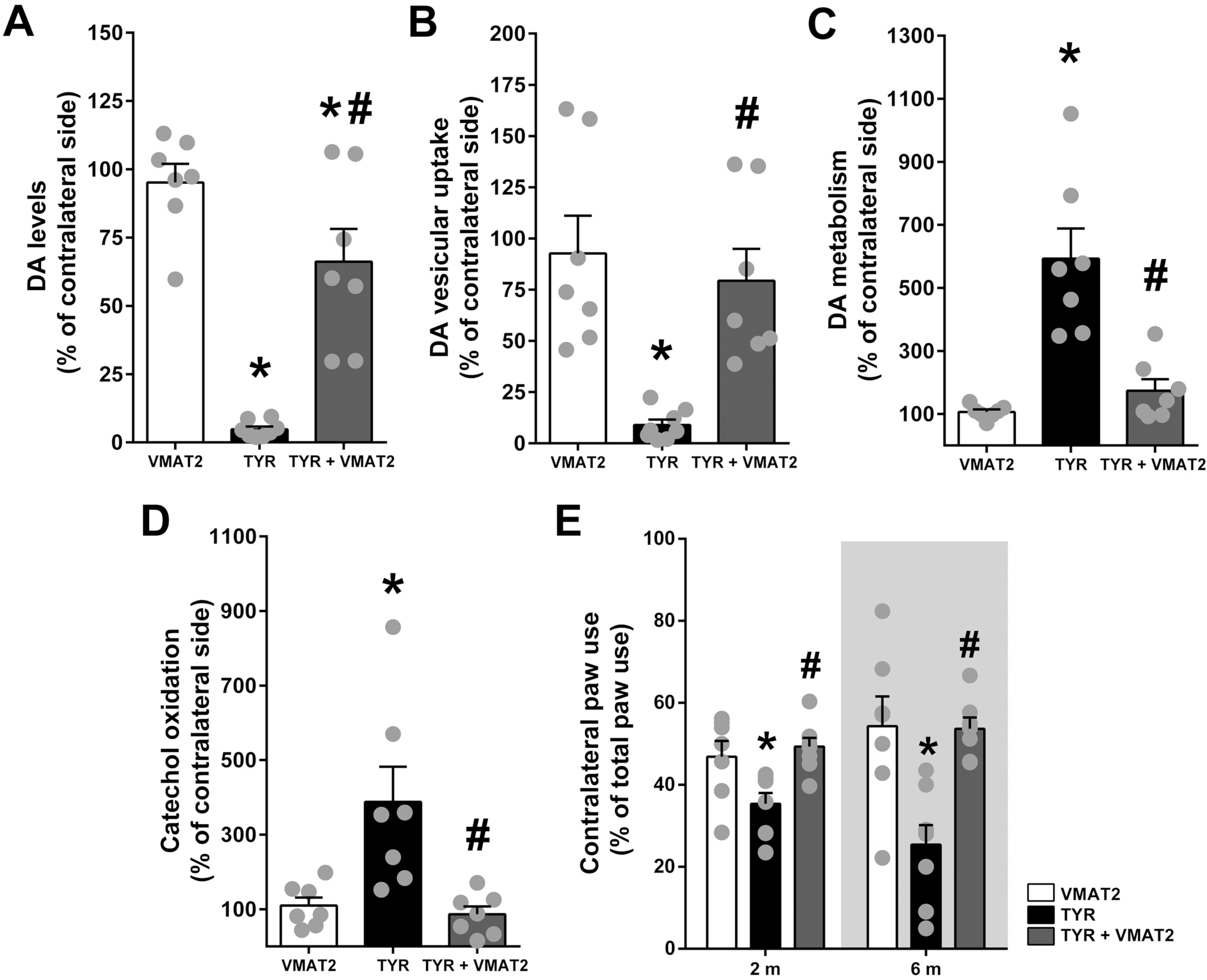
VMAT2 overexpression preserves striatal DA levels and motor function in NM-producing rats. UPLC-MS/MS quantification of striatal (**A**) DA levels; (**B**) DA vesicular uptake (calculated as DA:L-DOPA ratio); (**C**) DA metabolism (calculated as [3MT+DOPAC]:DA ratio); (**D**) catechol oxidation (calculated as [5SCD:L-DOPA]+[(5SCDA+AC):DA] ratio), in AAV-VMAT2, AAV-TYR and AAV-TYR+AAV-VMAT2-injected animals at 6m post-AAV injections. Results represent ipsilateral levels shown as the percentage of the contralateral (non-injected) side. (**E**) Contralateral forepaw use as assessed with the cylinder asymmetry test at 2m and 6m post-AAV injections. Values are mean ± SEM. \**p<*0.05 vs. VMAT2; ^#^*p<*0.05 vs. TYR, one-way ANOVA with Tukeýs post-hoc test. *N=* 7 (VMAT2), 7-8 (TYR), 7-8 (TYR+VMAT2) animals.

## Discussion

Modulation of VMAT2 activity has been in the spotlight as a potential therapeutic strategy for a wide range of diseases involving disturbances of DA metabolism and/or signaling, from addiction, schizophrenia or depression to neurodegenerative diseases such as Parkinson’s disease^6,30,31^. However, while some well-characterized pharmacological inhibitors of VMAT2 are available, such as reserpine and tetrabenazine, positive modulators of VMAT2 function are currently lacking^32^. Here, we found that viral vector-mediated VMAT2 overexpression provides therapeutic benefit in parkinsonian NM-producing rats by maintaining DA homeostasis, reducing potentially toxic DA oxidized species and preventing excessive age-dependent NM accumulation. The latter, in particular, represents the first demonstration that age-dependent NM production and subsequent intracellular buildup can be slowed down for therapeutic purposes *in vivo*. Indeed, until the recent introduction of the first rodent model of NM production^5^, NM had not been taken into account in experimental Parkinson’s disease *in vivo* modeling, despite the fact that this pigment has long been established as a major vulnerability factor for Parkinson’s disease-linked neurodegeneration^33^. This new melanized animal model revealed that continuous, age-dependent intracellular build-up of NM within autophagic structures ultimately leads to a general failure of cellular proteostasis associated to major Parkinson’s disease-like features, including motor deficits, LB-like inclusion formation and nigrostriatal neurodegeneration^5^. Here we show that all these pathological Parkinson’s disease-like features can be prevented or attenuated in these animals when lowering intracellular NM to levels below its pathogenic threshold by VMAT2 overexpression (see summary Supplementary Figure 4). Relevant to Parkinson’s disease, it has been reported in postmortem human brains that midbrain DA neurons with the highest VMAT2 protein expression exhibit the lowest NM levels and are the least vulnerable to Parkinson’s disease-linked neurodegeneration^13^. Conversely, the most vulnerable human midbrain DA neurons accumulate the most NM and have the lowest VMAT2 levels^13^.

Because VMAT2 overexpression reduces intracellular NM levels by decreasing free cytosolic DA that can subsequently oxidize into NM, the beneficial effects of VMAT2 could be related to reduced levels of potentially toxic oxidized DA species that serve as NM precursors. In particular, tyrosine, L-DOPA or DA-derived [o-]quinones, which are rapidly transformed into NM, have long been proposed as potential pathogenic factors in Parkinson’s disease^7^. In fact, NM synthesis is regarded as a protective antioxidant mechanism by trapping cytosolic quinones and semiquinones in lysosome-associated organelles so that they are no longer reactive with cytosolic components^34^. However, while high concentrations of DA can be acutely toxic *in vitro*^35^, the chronic enhancement of DA levels and oxidation does not cause DA nerve terminal or cell body loss *in vivo*^36–38^. For instance, chronic L-DOPA treatment^36,37^ or genetic enhancement of TH activity^38^ in rodents, both resulting in increased levels of DA and oxidized DA species, are not toxic to regular, wild-type animals but only to animals displaying additional Parkinson’s disease-related alterations, such as DJ-1 deficiency^36^ or A53T α-synuclein overexpression^38^. Relevant to humans, there is no clinical evidence for L-DOPA being neurotoxic and detrimental to the progression of Parkinson’s disease symptoms^39^. In addition, antioxidant strategies have systematically failed to provide any therapeutic benefit in Parkinson’s disease clinical trials^40^. Even if NM synthesis *per se* was initially neuroprotective, its long-term accumulation until occupying most of the neuronal cytoplasm has deleterious consequences by physically interfering with intracellular communication^41–43^ and impairing proteostasis^5,33^. In fact, L-DOPA seems to be toxic *in vitro* only at doses associated with NM formation^35^, which has been attributed to an interference by NM with intracellular neurotrophin signaling, and not to an acute toxic effect of L-DOPA *per se*^44^. Consistent with a macromolecular crowding effect linked to progressive NM accumulation, in-depth ultrastructural analyses have revealed that Lewy pathology in NM-containing neurons from Parkinson’s disease brains mostly consists of a crowded environment of vesicular structures and dysmorphic organelles^26^. In agreement with this, here we found that decreased intracellular NM levels by VMAT2 overexpression was associated to reduced Parkinson’s disease-like inclusion formation *in vivo*. In any case, any putative contribution of these species to NM-linked Parkinson’s disease pathology would be therapeutically targeted by VMAT2 overexpression, as the latter reduces both oxidized DA species and age-dependent NM accumulation.

The regulation of the balance of cytosolic DA levels is complex and involves an interplay between different processes. From one hand, dopamine transporter (DAT)-mediated reuptake of extracellular DA^45–47^, leaking from synaptic vesicles, and TH-mediated DA synthesis increase cytosolic DA levels. On the other hand, VMAT2-mediated DA encapsulation into synaptic vesicles and MAO-mediated DA metabolism decrease cytosolic DA levels^48^. While tyrosinase can mediate L-DOPA synthesis from tyrosine^48^ and thus accelerate DA production, the overall levels of cytosolic DA in TYR animals are not expected to remain significantly increased due to the feedback inhibition of cytosolic DA on TH^21,49^ and the transient condition of cytosolic DA, which is either rapidly metabolized or oxidized. In fact, in these animals TYR promotes the continuous oxidation of cytosolic DA into NM. In addition, the progressive accumulation of NM was associated to a marked downregulation of VMAT2 and TH that may contribute to the decreased DA vesicular uptake and vesicular DA levels^50^ observed in these animals. Furthermore, we have previously reported that striatal DAT levels are decreased by ∼70% in AAV-TYR-injected animals by 2 months post-AAV-TYR injections^5^, which concur with neuroimaging in vivo analyses in PD patients exhibiting reductions in DAT striatal density at early stages of the disease^51^. In this context, it is important to recall that DAT is also expressed in astrocytes and macrophages, which contribute to DA clearance from the synaptic cleft^46,47^. The increased number of CD68- and GFAP-positive cells in AAV-TYR-injected animals could thus favor an increased DA uptake and metabolism in glial cells, which is consistent with increased 3MT levels observed in the ventral midbrain of AAV-TYR-injected animals at 2 months post-AAV injection (Supplementary Table 4) and that could restrict the amount of DA available to be re-uptaken by neuronal DAT. All these changes in DA metabolism, combined with the progressive buildup of NM within the neuronal cytosol, may potentially contribute to the vulnerability of the dopaminergic system in TYR animals and in PD patients.

In addition to attenuate nigrostriatal neurodegeneration, VMAT2 overexpression also mitigated the phenotypic loss of TH expression that occurs within NM-laden neurons at early stages of degeneration. Indeed, despite the general notion that the number of DA neurons are reduced in Parkinson’s disease, NM-containing neurons from both Parkinson’s disease patients and parkinsonian NM-producing rats exhibit an early phenotypic loss of TH expression, associated to impaired DA release and early motor deficits, prior to degeneration^5,52^. In agreement with this, genetic disruption of mitochondrial complex I in mice has been recently reported to induce progressive parkinsonism in which TH downregulation and defective DA release in both striatal terminals and SNpc neuronal cell bodies precede neurodegeneration^53^. The occurrence of neuronal dysfunction before overt cell death and its restoration by VMAT2 may have important therapeutic implications, as it provides a therapeutic window in which neuronal function can be restored before the actual loss of the cell. The preservation of neuronal function by VMAT2 overexpression was not limited to a protection of nigrostriatal DA fibers and a maintenance of TH protein expression but also to a direct effect at maintaining DA homeostasis/metabolism, all of which resulted in an early and long-lasting preservation of the motor function in NM-producing parkinsonian rats. Importantly, overexpression of VMAT2 by itself did not alter the normal motor function in non-parkinsonian (control) animals. This is in agreement with previous studies in VMAT2-overexpressing transgenic mice^54^. Indeed, while VMAT2 overexpression has been reported to increase DA quantal size and DA release^55^, which could theoretically result in behavioral changes, VMAT2 overexpression in transgenic mice did not result in any dramatic behavioral phenotype, as measured by motor, sensory, affective, appetitive, or social assays^54^. From a translational point of view, these observations support the feasibility and safety of increasing VMAT2 function as a potential therapeutic strategy in humans.

Overall, our results demonstrate for the first time the feasibility and therapeutic potential of modulating NM production *in vivo* by modifying DA homeostasis/metabolism with VMAT2. These results open the possibility of targeting NM production to prevent or delay Parkinson’s disease by slowing down the progressive accumulation of NM that occurs with age. Such a strategy would represent a conceptually novel therapeutic approach for Parkinson’s disease and, in a broader sense, brain aging. There are, however, some limitations of this study. For instance, by concomitantly injecting AAV-VMAT2 and AAV-TYR, the production of NM was slowed down from its initial synthesis, thus being able to prevent or delay disease onset. It remains to be determined whether in a Parkinson’s disease patient in which intracellular NM has already reached pathological levels, VMAT2 would be able to halt or delay disease progression. Finally, while we have shown a beneficial effect of VMAT2 overexpression up to 6 months post-AAV injection, we do not know how much longer this beneficial effect would potentially last. Indeed, VMAT2 overexpression may slow down but not completely halt NM production, so NM could continue to steadily accumulate with age despite VMAT2 overexpression, although at a slower rate. To overcome this limitation, VMAT2 overexpression could be potentially combined with strategies aimed at eliminating intracellular NM once it has already been produced, such as by overexpression of autophagy master regulator TFEB, which we have recently shown to promote the clearance of NM-filled autophagic structures^56^.

## Acknowledgements

We are grateful to Dr. Iria Carballo-Carbajal for helping prepare the AAV-VMAT2, to Professor Kazumasa Wakamatsu and Professor Shosuke Ito (Fujita Health University, Aichi, Japan) for providing both SCD and 5SCDA standards and to the U20/FVPR unit of ICTS-NANBIOSIS at the VHIR (Barcelona, Spain) for assistance in histological processing.

## Supplementary material

Supplementary material is available at *Brain* online.

---

**Supplementary Figure 1.**
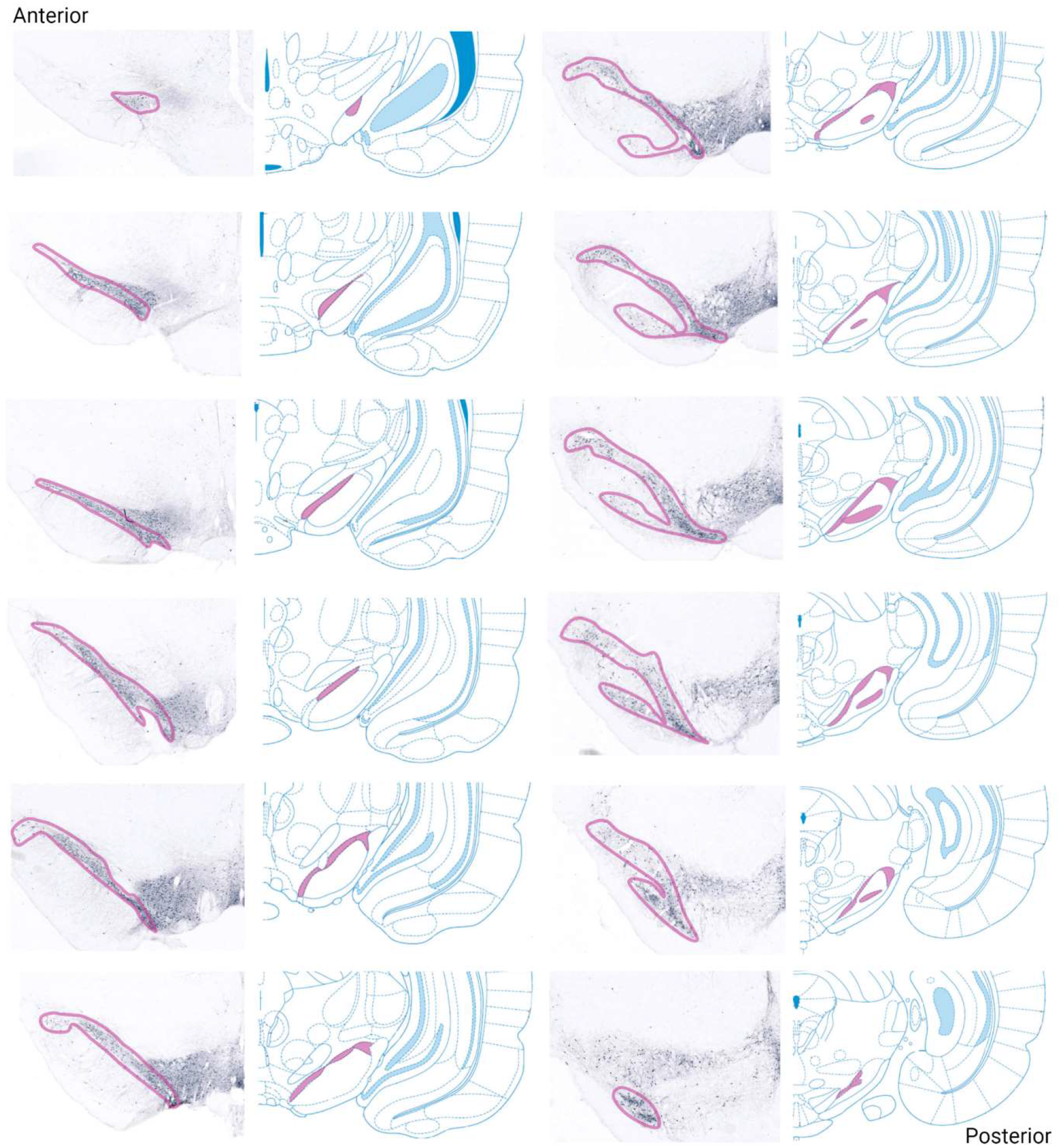
Anatomical levels of rat SN used for quantifications. Twelve serial 5 µm-thick paraffin sections covering the entire rat SNpc (one every 17^th^ section) were immunostained for TH and matched with the corresponding anatomical level from the rat brain atlas (Paxinos and Watson, 2004). Pink color indicates the borders of the SNpc in both the atlas and TH-immunostained sections. Figure was created with BioRender.com.

**Supplementary Figure 2.**
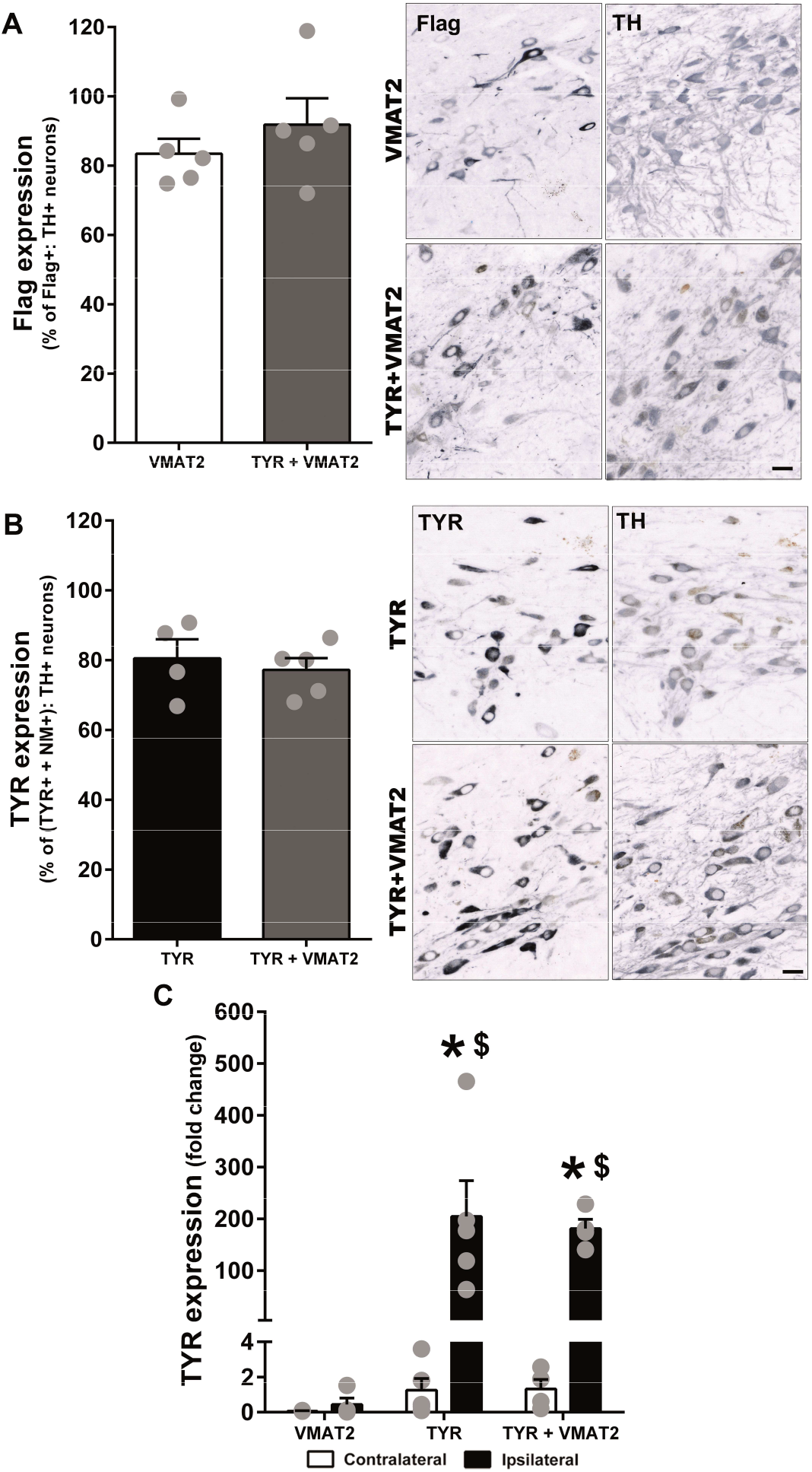
TYR and VMAT2 transduction efficiency in AAV-TYR- and AAV-VMAT2-injected animals, alone or in combination. **(A-B)** Percentage of SNpc TH-positive neurons expressing Flag-VMAT2 **(A)** or TYR **(B)** at 1m post-AAV injections, as assessed in serial 5 µm-thick paraffin adjacent sections covering the entire SN (every 17th section for a total of 10-12 sections analyzed/animal). Photomicrographs correspond to representative images of adjacent SNpc sections immunostained for either **(A)** Flag or TH (in gray, unstained NM in brown) and **(B)** TYR or TH (in gray, unstained NM in brown). Scale bars: 20 µm. **(C)** TYR mRNA levels assessed by qPCR in contralateral and ipsilateral vMB samples from VMAT, TYR and TYR+VMAT2 animals. In all panels, values are mean±SEM. In **(A-B)**, N.S. (two-tailed t-test). In **(C)**, \**p*<0.05 vs. ipsilateral VMAT2; ^$^p<0.05 vs. respective contralateral side; two-way ANOVA with Tukeýs post-hoc test. In **(A-B)** N= 5 (VMAT2), 4 (TYR), 5 (TYR+VMAT2) animals. In **(C)** N= 3-4 (VMAT2), 5 (TYR), 4 (TYR+VMAT2) animals.

**Supplementary Figure 3.**
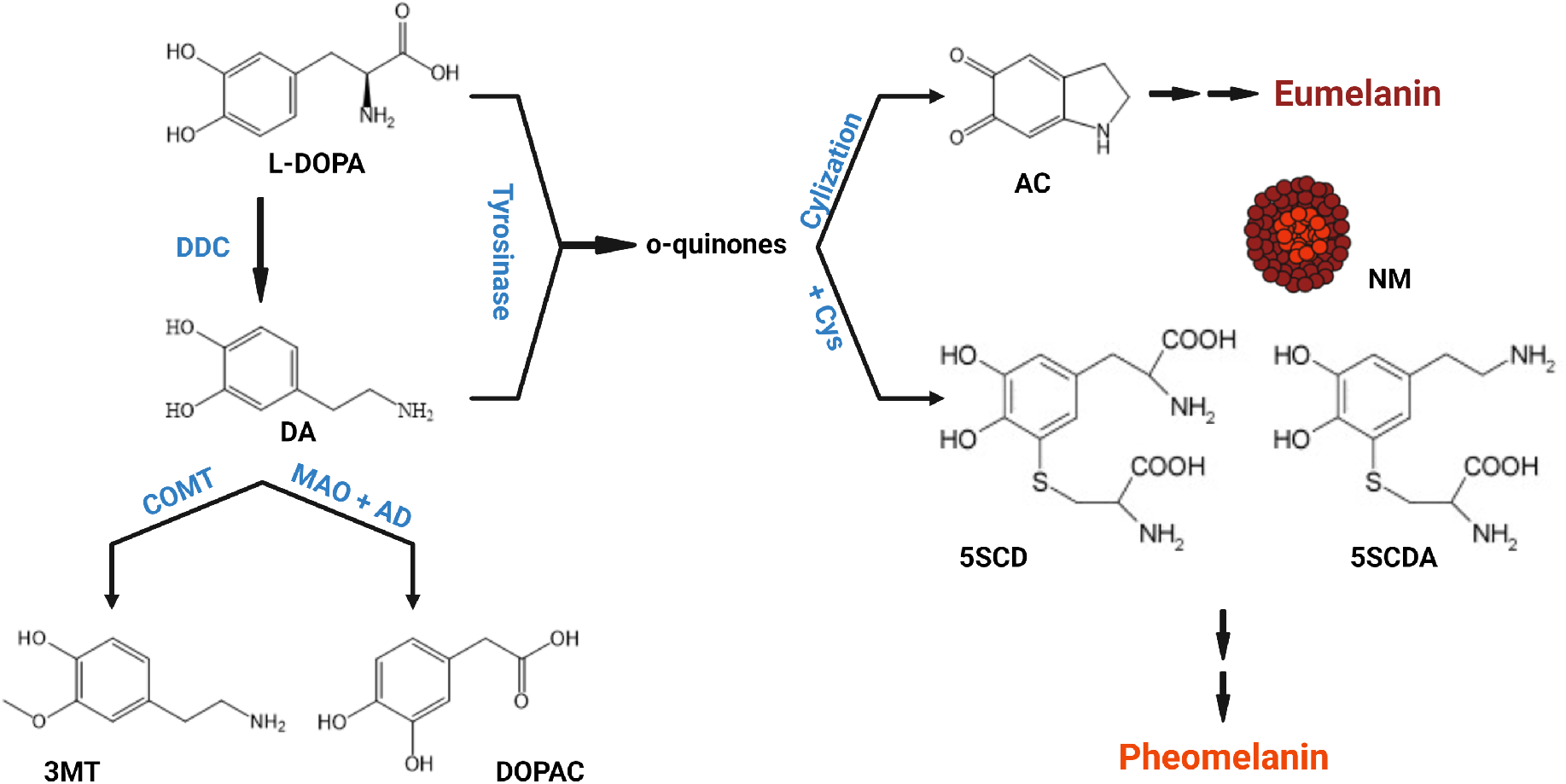
Schematic representation of DA metabolism and oxidation pathways. DA is synthesized in cytoplasm from L-DOPA by DDC. Cytosolic DA can be degraded by MAO to produce 3MT or by MAO followed by AD to produce DOPAC. On the other hand, both cytosolic DA and L-DOPA can be oxidized either spontaneously or by tyrosinase to produce o-quinones. These, in turn generate AC, 5SCDA and 5SCD which will act as precursors of the eumelanin and pheomelanin components of NM. 5SCD, 5-*S*-cysteinyldopa; 5SCDA, 5-*S*-cysteinyldopamine; AC, aminochrome; AD, aldehyde dehydrogenase; COMT, catechol-O-methyltransferase; DA, dopamine; DDC, dopa decarboxylase; DOPAC, 3,4-dihydroxyphenylacetic acid; L-DOPA, 3,4-dihydroxyphenylalanine; MAO, monoamine oxidase; 3-MT, 3-methoxytyramine; NM, neuromelanin. Figure was created with BioRender.com.

**Supplementary Figure 4.**
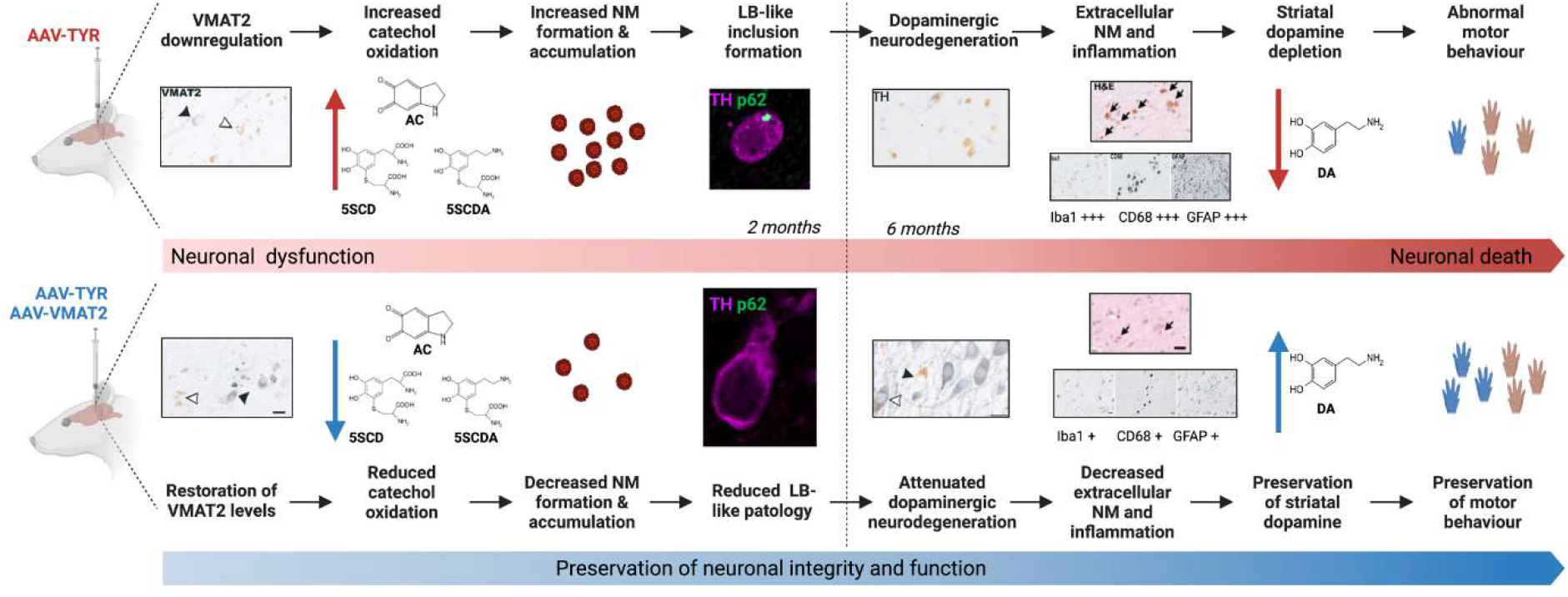
Graphical summary of the main results. See main text for details. Figure was created with BioRender.com.

**Supplementary Table 1.** UPLC-MS/MS assessment of striatal DA and L-DOPA levels (pmol/mg ± SEM) and DA vesicular uptake ratio in AAV-VMAT2 injected rats at 2 months (m) post-injection.

| Group | Hemisphere | Animal | DA | L-DOPA | DA vesicular uptake (DA/L-DOPA) |
| --- | --- | --- | --- | --- | --- |
| AAV-VMAT2 | Contralateral | 1 | 189,91 | 0,96 | 196,94 |
|  |  | 2 | 254,86 | 2,02 | 126,21 |
|  |  | 3 | 144,06 | 3,54 | 40,72 |
|  |  | 4 | 176,18 | 1,34 | 131,06 |
|  |  | 5 | 326,49 | 5,48 | 59,56 |
|  |  | 6 | 313,83 | 6,93 | 45,28 |
|  |  | 7 | 245,14 | 4,42 | 55,48 |
|  |  | 8 | 313,61 | 8,69 | 36,08 |
|  | Ipsilateral | 1 | 244,76 | 1,43 | 171,47 |
|  |  | 2 | 322,61 | 1,14 | 282,03 |
|  |  | 3 | 250,39 | 1,23 | 204,31 |
|  |  | 4 | 300,48 | 2,14 | 140,35 |
|  |  | 5 | 313,82 | 0,93 | 337,73 |
|  |  | 6 | 286,01 | 1,26 | 226,60 |
|  |  | 7 | 384,66 | 3,16 | 121,59 |
|  |  | 8 | 221,20 | 1,12 | 197,14 |

**Supplementary Table 2.** Cell counts in rat SNpc at 1m post-AAV injections.

|  | Hemisphere | Animal | Flag expression |  |  |  | TYR expression |  |  |  |
| --- | --- | --- | --- | --- | --- | --- | --- | --- | --- | --- |
|  |  |  | Flag+ | Flag-NM+ | TH+ | TH-NM+ | TYR+ | TYR-NM+ | TH+ | TH-NM+ |
| **AAV-VMAT2** | **Ipsilateral** | 1 | 241 | N/A | 286 | N/A | N/A | N/A | N/A | N/A |
|  |  | 2 | 460 | N/A | 560 | N/A | N/A | N/A | N/A | N/A |
|  |  | 3 | 170 | N/A | 227 | N/A | N/A | N/A | N/A | N/A |
|  |  | 4 | 538 | N/A | 703 | N/A | N/A | N/A | N/A | N/A |
|  |  | 5 | 395 | N/A | 398 | N/A | N/A | N/A | N/A | N/A |
| **AAV-TYR** | **Ipsilateral** | 1 | N/A | N/A | N/A | N/A | 1054 | 9 | 836 | 15 |
|  |  | 2 | N/A | N/A | N/A | N/A | 1152 | 18 | 861 | 3 |
|  |  | 3 | N/A | N/A | N/A | N/A | 765 | 7 | 783 | 2 |
|  |  | 4 | N/A | N/A | N/A | N/A | 787 | 9 | 809 | 1 |
| **AAV-TYR +**<br>**AAV-VMAT2** | **Ipsilateral** | 1 | 643 | 14 | 754 | 6 | 706 | 59 | 860 | 1 |
|  |  | 2 | 592 | 6 | 822 | 8 | 650 | 27 | 589 | 8 |
|  |  | 3 | 656 | 18 | 747 | 1 | 555 | 40 | 747 | 3 |
|  |  | 4 | 723 | 2 | 789 | 2 | 580 | 40 | 734 | 1 |
|  |  | 5 | 804 | 2 | 675 | 3 | 863 | 13 | 779 | 2 |

N/A not aplicable

**Supplementary Table 3.** Cell counts in rat SNpc at 2m post-AAV injections.

|  | Hemisphere | Animal | VMAT2 |  |  | p62+ Inclusions |  |
| --- | --- | --- | --- | --- | --- | --- | --- |
|  |  |  | VMAT2+<br>NM- | VMAT2+<br>NM+ | VMAT2-<br>NM+ | cytoplasmic | nuclear |
| AAV-VMAT2 | Contralateral | 1 | 51 | N/A | N/A | N/A | N/A |
|  |  | 2 | 46 | N/A | N/A | N/A | N/A |
|  |  | 3 | 58 | N/A | N/A | N/A | N/A |
|  |  | 4 | 46 | N/A | N/A | N/A | N/A |
|  |  | 5 | 74 | N/A | N/A | N/A | N/A |
|  |  | 6 | 67 | N/A | N/A | N/A | N/A |
|  |  | 7 | 48 | N/A | N/A | N/A | N/A |
|  | Ipsilateral | 1 | 64 | N/A | N/A | N/A | N/A |
|  |  | 2 | 64 | N/A | N/A | N/A | N/A |
|  |  | 3 | 93 | N/A | N/A | N/A | N/A |
|  |  | 4 | 33 | N/A | N/A | N/A | N/A |
|  |  | 5 | 34 | N/A | N/A | N/A | N/A |
|  |  | 6 | 60 | N/A | N/A | N/A | N/A |
|  |  | 7 | 63 | N/A | N/A | N/A | N/A |
| AAV-TYR | Contralateral | 1 | 51 | N/A | N/A | 0 | 0 |
|  |  | 2 | 33 | N/A | N/A | 0 | 0 |
|  |  | 3 | 45 | N/A | N/A | 0 | 0 |
|  |  | 4 | 90 | N/A | N/A | 0 | 0 |
|  |  | 5 | a | N/A | N/A | 0 | 0 |
|  |  | 6 | a | N/A | N/A | 0 | 0 |
|  |  | 7 | a | N/A | N/A | 0 | 0 |
|  |  | 8 | 75 | N/A | N/A | 0 | 0 |
|  | Ipsilateral | 1 | 0 | 3 | 42 | 6 | 0 |
|  |  | 2 | 12 | 45 | 48 | 0 | 0 |
|  |  | 3 | 0 | 3 | 147 | 5 | 0 |
|  |  | 4 | 0 | 21 | 90 | 2 | 0 |
|  |  | 5 | a | a | a | 5 | 0 |
|  |  | 6 | a | a | a | 3 | 0 |
|  |  | 7 | a | a | a | 1 | 0 |
|  |  | 8 | 0 | 33 | 54 | 4 | 0 |
| AAV-TYR +<br>AAV-VMAT2 | Contralateral | 1 | 39 | N/A | N/A | 0 | 0 |
|  |  | 2 | 42 | N/A | N/A | 0 | 0 |
|  |  | 3 | 51 | N/A | N/A | 0 | 0 |
|  |  | 4 | 78 | N/A | N/A | 0 | 0 |
|  |  | 5 | 30 | N/A | N/A | 0 | 0 |
|  |  | 6 | 66 | N/A | N/A | 0 | 0 |
|  |  | 7 | 69 | N/A | N/A | 0 | 0 |
|  |  | 8 | 36 | N/A | N/A | 0 | 0 |
|  | Ipsilateral | 1 | 6 | 6 | 15 | 1 | 0 |
|  |  | 2 | 30 | 21 | 18 | 0 | 0 |
|  |  | 3 | 42 | 21 | 39 | 0 | 0 |
|  |  | 4 | 15 | 9 | 21 | 0 | 0 |
|  |  | 5 | 18 | 33 | 27 | 2 | 0 |
|  |  | 6 | 21 | 18 | 9 | 0 | 0 |
|  |  | 7 | 33 | 30 | 33 | 0 | 0 |
|  |  | 8 | 6 | 9 | 12 | 0 | 0 |

a sections not available N/A not applicable

**Supplementary Table 4.**
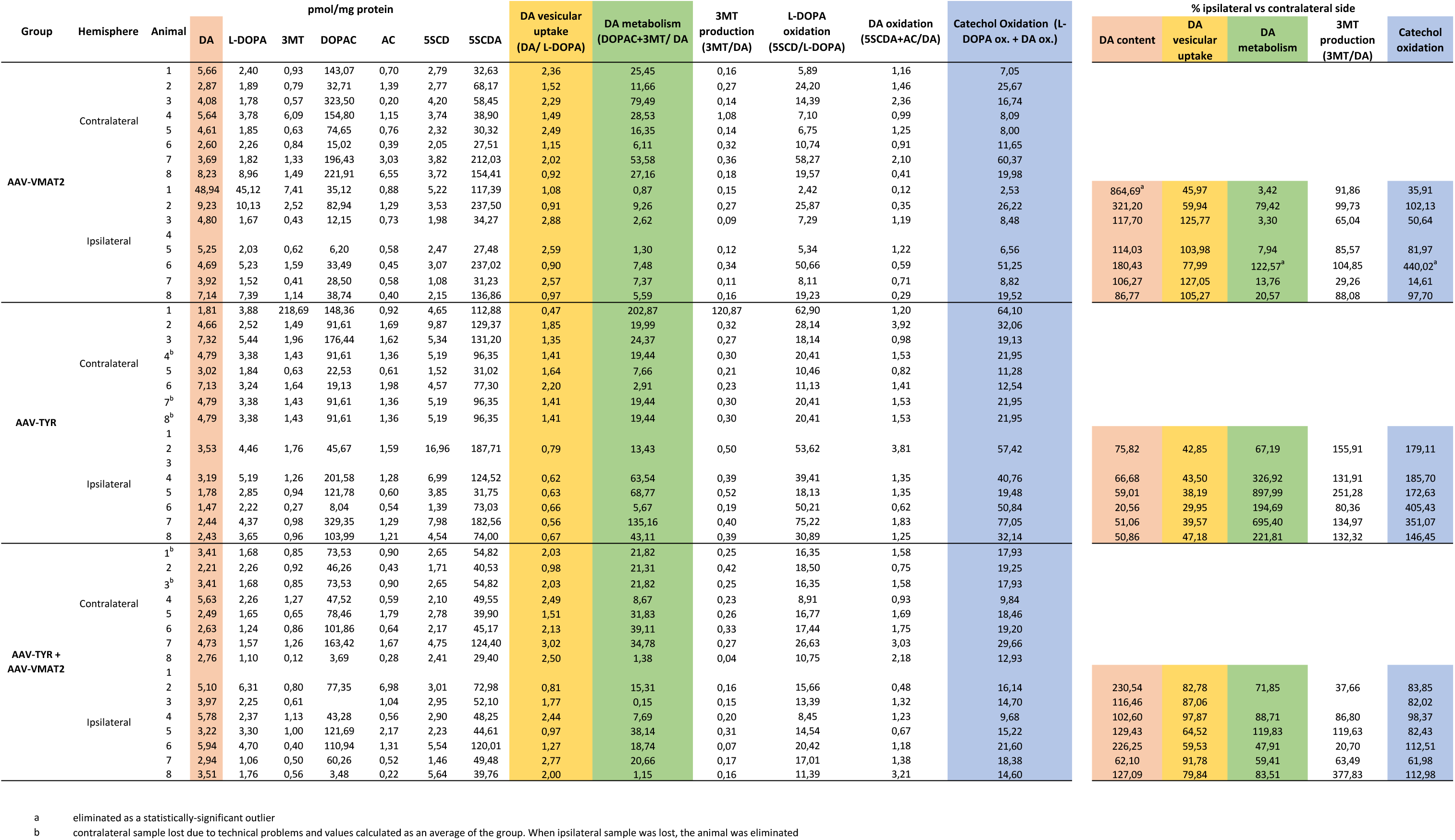
UPLC-MS/MS assessment of DA metabolites (pmol/mg ± SEM) and their calculated ratios (% ipsilateral vs contralateral) in rat ventral midbrain at 2m post-AAV injections.

**Supplementary Table 5.** UPLC-MS/MS assessment of DA metabolites (pmol/mg ± SEM) and their calculated ratios (% ipsilateral vs contralateral) in rat striatum at 2m post-AAV injections.

| Group | Hemisphere | Animal | pmol/mg protein |  |  |  |  |  |  | DA vesicular uptake (DA/ L-DOPA) | DA metabolism (DOPAC+3MT/ DA) | L-DOPA oxidation (5SCD/L-DOPA) | DA oxidation (5SCDA+AC/DA) | Catechol Oxidation (L-DOPA ox. + DA ox.) | % ipsilateral vs contralateral side |  |  |  |
| --- | --- | --- | --- | --- | --- | --- | --- | --- | --- | --- | --- | --- | --- | --- | --- | --- | --- | --- |
|  |  |  | DA | L-DOPA | 3MT | DOPAC | AC | 5SCD | 5SCDA |  |  |  |  |  | DA content | DA vesicular uptake | DA metabolism | Catechol oxidation |
| AAV-VMAT2 | Contralateral | 1 | 305,83 | 4,53 | 37,39 | 515,07 | 1,62 | 10,31 | 61,93 | 67,44 | 1,81 | 2,27 | 0,21 | 2,48 |  |  |  |  |
|  |  | 2 | 184,14 | 3,17 | 28,92 | 522,93 | 1,38 | 17,97 | 54,20 | 58,04 | 3,00 | 5,66 | 0,30 | 5,97 |  |  |  |  |
|  |  | 3 | 302,11 | 9,07 | 47,38 | 572,31 | 1,30 | 13,38 | 70,09 | 33,29 | 2,05 | 1,47 | 0,24 | 1,71 |  |  |  |  |
|  |  | 4 | 274,23 | 8,26 | 40,95 | 232,81 | 0,99 | 15,85 | 75,53 | 33,19 | 1,00 | 1,92 | 0,28 | 2,20 |  |  |  |  |
|  |  | 5 | 289,97 | 10,16 | 29,27 | 290,63 | 2,76 | 9,21 | 83,09 | 28,55 | 1,10 | 0,91 | 0,30 | 1,20 |  |  |  |  |
|  |  | 6 | 179,08 | 4,70 | 33,72 | 319,67 | 0,97 | 22,31 | 68,11 | 38,14 | 1,97 | 4,75 | 0,39 | 5,14 |  |  |  |  |
|  |  | 7 | 358,68 | 9,51 | 47,53 | 326,00 | 2,89 | 13,09 | 61,69 | 37,74 | 1,04 | 1,38 | 0,18 | 1,56 |  |  |  |  |
|  |  | 8 | 335,04 | 8,82 | 46,83 | 685,21 | 0,71 | 15,23 | 84,59 | 37,98 | 2,18 | 1,73 | 0,25 | 1,98 |  |  |  |  |
|  | Ipsilateral | 1 | 249,31 | 5,32 | 50,04 | 263,62 | 1,05 | 12,85 | 62,96 | 46,82 | 1,26 | 2,41 | 0,26 | 2,67 | 81,52 | 69,42 | 69,64 | 107,63 |
|  |  | 2 | 213,21 | 7,14 | 31,11 | 202,78 | 0,99 | 14,72 | 51,25 | 29,86 | 1,10 | 2,06 | 0,25 | 2,31 | 115,79 | 51,45 | 36,61 | 38,66 |
|  |  | 3 | 304,50 | 8,46 | 35,45 | 263,63 | 0,35 | 12,21 | 37,24 | 35,99 | 0,98 | 1,44 | 0,12 | 1,57 | 100,79 | 108,10 | 47,88 | 91,57 |
|  |  | 4 | 270,00 | 6,25 | 50,16 | 191,11 | 1,16 | 20,38 | 61,02 | 43,18 | 0,89 | 3,26 | 0,23 | 3,49 | 98,46 | 130,12 | 89,52 | 158,89 |
|  |  | 5 | 333,18 | 14,76 | 38,16 | 362,66 | 2,24 | 11,20 | 98,77 | 22,57 | 1,20 | 0,76 | 0,30 | 1,06 | 114,90 | 79,04 | 109,04 | 88,28 |
|  |  | 6 | 274,98 | 7,61 | 39,47 | 375,71 | 1,34 | 18,58 | 67,04 | 36,16 | 1,51 | 2,44 | 0,25 | 2,69 | 153,55 | 94,81 | 76,51 | 52,39 |
|  |  | 7 | 297,05 | 36,84 | 33,86 | 216,22 | 4,33 | 9,80 | 58,42 | 8,06 | 0,84 | 0,27 | 0,21 | 0,48 | 82,82 | 21,37 | 80,84 | 30,65 |
|  |  | 8 | 225,98 | 11,07 | 24,81 | 203,06 | 1,33 | 4,56 | 96,69 | 20,41 | 1,01 | 0,41 | 0,43 | 0,85 | 67,45 | 53,73 | 46,15 | 42,70 |
| AAV-TYR | Contralateral | 1 | 223,28 | 4,79 | 26,13 | 140,39 | 1,71 | 17,49 | 410,40 | 46,65 | 0,75 | 3,65 | 1,85 | 5,50 |  |  |  |  |
|  |  | 2 | 370,58 | 11,28 | 32,22 | 259,43 | 2,58 | 9,92 | 96,46 | 32,85 | 0,79 | 0,88 | 0,27 | 1,15 |  |  |  |  |
|  |  | 3 | 363,22 | 5,97 | 26,80 | 258,69 | 2,20 | 5,61 | 223,17 | 60,88 | 0,79 | 0,94 | 0,62 | 1,56 |  |  |  |  |
|  |  | 4 | 170,31 | 4,80 | 18,23 | 200,02 | 0,24 | 22,32 | 193,56 | 35,46 | 1,28 | 4,65 | 1,14 | 5,78 |  |  |  |  |
|  |  | 5 | 190,06 | 13,81 | 22,55 | 214,02 | 3,72 | 16,96 | 212,61 | 13,77 | 1,24 | 1,23 | 1,14 | 2,37 |  |  |  |  |
|  |  | 6 | 183,94 | 6,07 | 22,06 | 194,21 | 2,11 | 26,18 | 157,79 | 30,33 | 1,18 | 4,32 | 0,87 | 5,19 |  |  |  |  |
|  |  | 7 | 363,57 | 9,55 | 42,05 | 167,76 | 2,81 | 20,65 | 224,67 | 38,08 | 0,58 | 2,16 | 0,63 | 2,79 |  |  |  |  |
|  |  | 8 | 316,20 | 14,41 | 29,72 | 256,50 | 3,98 | 13,58 | 262,20 | 21,94 | 0,91 | 0,94 | 0,84 | 1,78 |  |  |  |  |
|  | Ipsilateral | 1 | 36,55 | 3,52 | 8,26 | 27,88 | 0,77 | 20,39 | 187,67 | 10,39 | 0,99 | 5,80 | 5,16 | 10,95 | 16,37 | 22,28 | 132,56 | 199,18 |
|  |  | 2 | 40,26 | 6,44 | 14,60 | 46,65 | 0,53 | 45,11 | 151,12 | 6,25 | 1,52 | 7,00 | 3,77 | 10,77 | 10,86 | 19,02 | 193,31 | 939,32<sup>a</sup> |
|  |  | 3 | 44,06 | 10,11 | 6,53 | 158,16 | 1,18 | 15,58 | 126,29 | 4,36 | 3,74 | 1,54 | 2,89 | 4,43 | 12,13 | 7,16 | 475,49 | 284,01 |
|  |  | 4 | 42,19 | 5,60 | 7,32 | 43,42 | 0,56 | 61,58 | 260,62 | 7,54 | 1,20 | 11,01 | 6,19 | 17,20 | 24,77 | 21,26 | 93,86 | 297,27 |
|  |  | 5 | 216,45 | 7,30 | 19,93 | 191,24 | 2,49 | 21,22 | 1140,05 | 29,66 | 0,98 | 2,91 | 5,28 | 8,19 | 113,89<sup>a</sup> | 215,48<sup>a</sup> | 78,38 | 345,89 |
|  |  | 6 | 48,23 | 6,65 | 7,24 | 118,12 | 2,72 | 20,46 | 194,05 | 7,25 | 2,60 | 3,08 | 4,08 | 7,16 | 26,22 | 23,92 | 221,09 | 138,03 |
|  |  | 7 | 66,50 | 10,64 | 10,84 | 137,43 | 3,88 | 30,36 | 155,48 | 6,25 | 2,23 | 2,85 | 2,40 | 5,25 | 18,29 | 16,42 | 386,34 | 188,32 |
|  |  | 8 | 87,61 | 12,90 | 11,33 | 93,64 | 2,46 | 23,10 | 152,97 | 6,79 | 1,20 | 1,79 | 1,77 | 3,56 | 27,71 | 30,95 | 132,36 | 199,76 |
| AAV-TYR + AAV-VMAT2 | Contralateral | 1 | 360,23 | 8,25 | 62,93 | 500,16 | 1,60 | 21,24 | 146,80 | 43,65 | 1,56 | 2,57 | 0,41 | 2,99 |  |  |  |  |
|  |  | 2 | 280,94 | 8,17 | 31,82 | 278,06 | 1,43 | 7,14 | 55,22 | 34,39 | 1,10 | 0,87 | 0,20 | 1,08 |  |  |  |  |
|  |  | 3 | 276,02 | 4,58 | 45,97 | 400,51 | 1,33 | 29,70 | 219,27 | 60,24 | 1,62 | 6,48 | 0,80 | 7,28 |  |  |  |  |
|  |  | 4 | 162,19 | 3,87 | 22,99 | 203,33 | 0,41 | 19,52 | 61,04 | 41,94 | 1,40 | 5,05 | 0,38 | 5,43 |  |  |  |  |
|  |  | 5 | 365,05 | 10,10 | 39,52 | 195,26 | 1,41 | 30,70 | 381,08 | 36,16 | 0,64 | 3,04 | 1,05 | 4,09 |  |  |  |  |
|  |  | 6 | 38,72 | 5,72 | 8,68 | 89,57 | 1,20 | 7,62 | 67,64 | 6,77 | 2,54 | 1,33 | 1,78 | 3,11 |  |  |  |  |
|  |  | 7 | 295,69 | 8,37 | 34,22 | 152,04 | 0,67 | 24,42 | 547,94 | 35,31 | 0,63 | 2,92 | 1,86 | 4,77 |  |  |  |  |
|  |  | 8 | 220,17 | 11,11 | 27,72 | 224,97 | 1,26 | 10,98 | 64,55 | 19,81 | 1,15 | 0,99 | 0,30 | 1,29 |  |  |  |  |
|  | Ipsilateral | 1 | 214,02 | 6,49 | 32,44 | 355,03 | 0,87 | 21,45 | 58,20 | 32,98 | 1,81 | 3,31 | 0,28 | 3,58 | 59,41 | 75,56 | 115,82 | 119,95 |
|  |  | 2 | 136,11 | 34,43 | 20,27 | 130,68 | 1,02 | 13,85 | 106,60 | 3,95 | 1,11 | 0,40 | 0,79 | 1,19 | 48,45 | 11,50 | 100,55 | 110,89 |
|  |  | 3 | 202,51 | 6,47 | 34,93 | 129,91 | 0,95 | 20,67 | 105,46 | 31,31 | 0,81 | 3,20 | 0,53 | 3,72 | 73,37 | 51,97 | 50,32 | 51,10 |
|  |  | 4 | 270,38 | 8,23 | 18,69 | 165,19 | 1,07 | 11,77 | 199,31 | 32,85 | 0,68 | 1,43 | 0,74 | 2,17 | 166,70 | 78,32 | 48,74 | 40,01 |
|  |  | 5 | 80,44 | 3,30 | 16,35 | 39,55 | 0,55 | 25,98 | 150,70 | 24,39 | 0,69 | 7,88 | 1,88 | 9,76 | 22,04 | 67,45 | 108,05 | 238,64 |
|  |  | 6 | 85,29 | 9,78 | 14,00 | 136,41 | 1,37 | 15,09 | 132,26 | 8,72 | 1,76 | 1,54 | 1,57 | 3,11 | 220,28 | 128,82 | 69,49 | 99,96 |
|  |  | 7 | 162,35 | 10,27 | 28,52 | 88,88 | 0,89 | 31,53 | 361,25 | 15,81 | 0,72 | 3,07 | 2,23 | 5,30 | 54,91 | 44,79 | 114,80 | 111,12 |
|  |  | 8 | 115,35 | 11,97 | 18,02 | 86,24 | 0,70 | 21,50 | 252,73 | 9,64 | 0,90 | 1,80 | 2,20 | 3,99 | 52,39 | 48,64 | 78,76 | 310,20 |

a eliminated as a statistically-significant outlier

**Supplementary Table 6.**
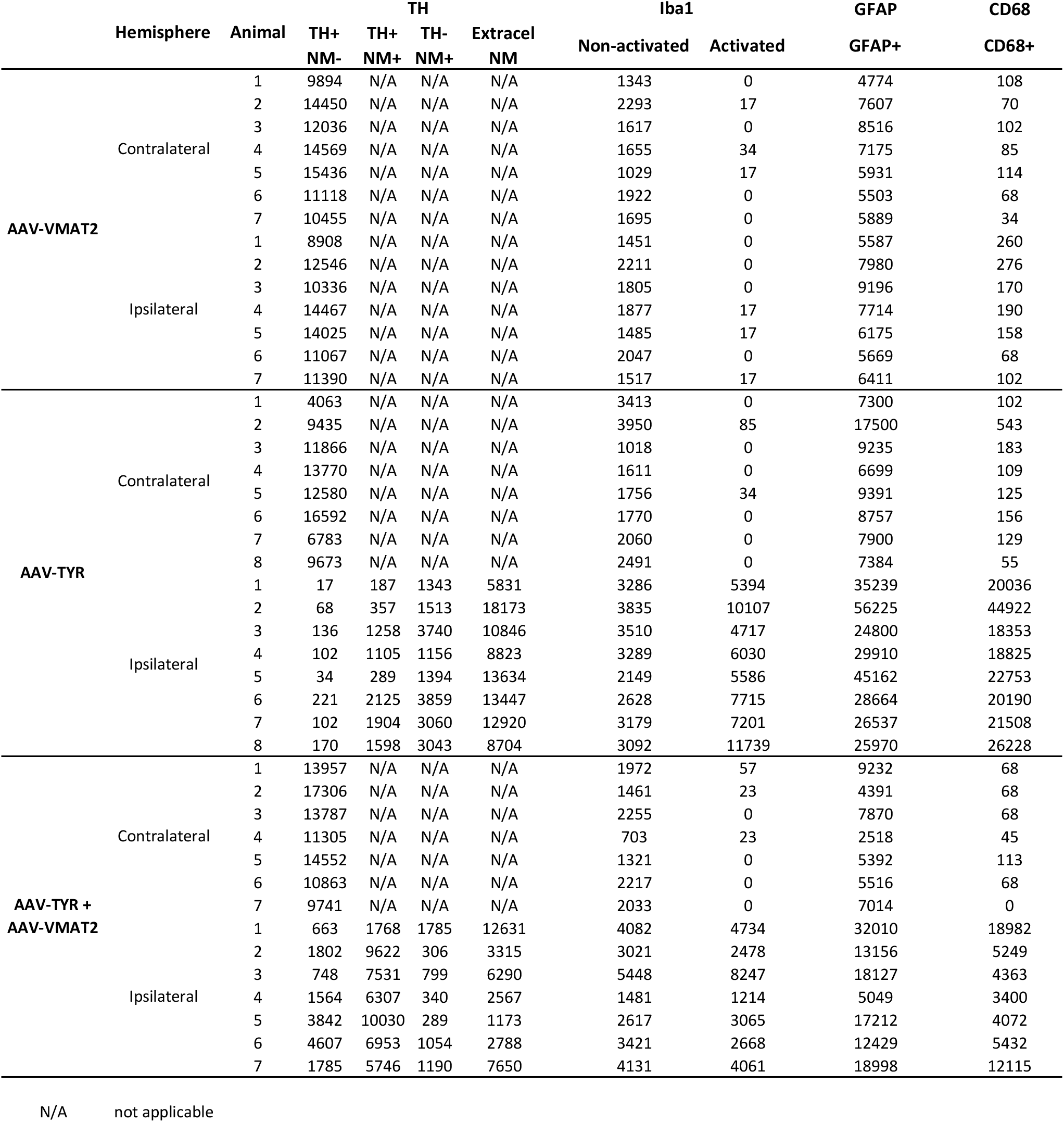
Cell counts in rat SNpc at 6m post-AAV injections.

**Supplementary Table 7.**
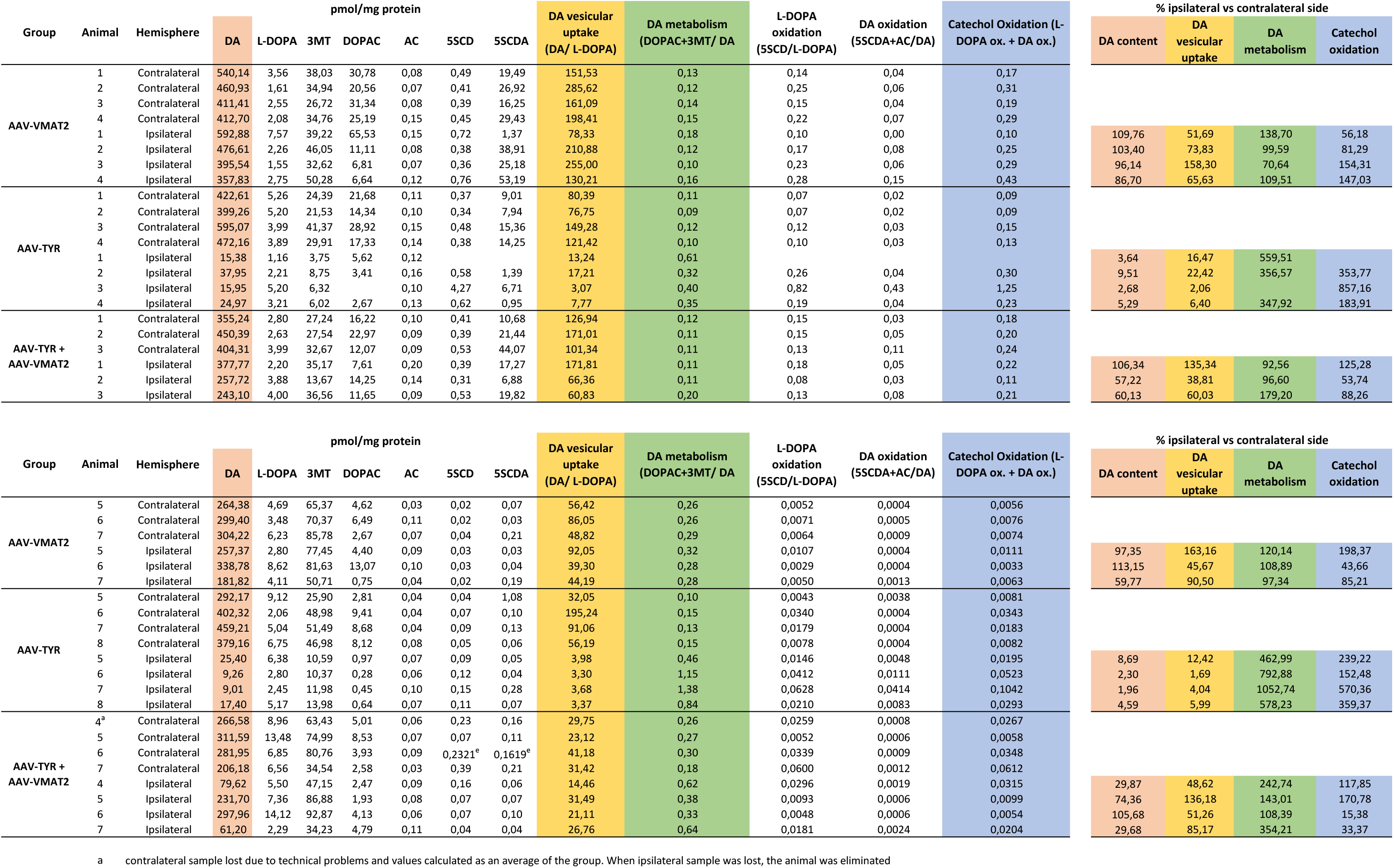
UPLC-MS/MS assessment of DA metabolites (pmol/mg ± SEM) and their calculated ratios (% ipsilateral vs contralateral) in rat striatum at 6m post-AAV injections.

